# The Induction of Pyrenoid Synthesis by Hyperoxia and its Implications for the Natural Diversity of Photosynthetic Responses in *Chlamydomonas*

**DOI:** 10.1101/2021.03.10.434646

**Authors:** Peter Neofotis, Joshua Temple, Oliver L. Tessmer, Jacob Bibik, Nicole Norris, Eric Poliner, Ben Lucker, Sarathi Wijetilleke, Alecia Withrow, Barbara Sears, Greg Mogos, Melinda Frame, David Hall, Joseph Weissman, David M. Kramer

**Affiliations:** MSU-DOE Plant Research Laboratory, Michigan State University, East Lansing, MI; Center for Advanced Microscopy, Michigan State University, East Lansing, MI; Great Lakes Bioenergy Research Center, Michigan State University, East Lansing, MI; Corporate Strategic Research, ExxonMobil, Annandale, NJ

## Abstract

In algae, it is well established that the pyrenoid, a component of the carbon-concentrating mechanism (CCM), is essential for efficient photosynthesis at low CO_2_. However, the signal that triggers the formation of the pyrenoid has remained elusive. Here, we show that, in *Chlamydomonas reinhardtii*, the pyrenoid is strongly induced by hyperoxia, even at high CO_2_ or bicarbonate levels. These results suggest that the pyrenoid can be induced by a common product of photosynthesis specific to low CO_2_ or hyperoxia. Consistent with this view, the photorespiratory by-product, H_2_O_2_, induced the pyrenoid, suggesting that it acts as a signal. Finally, we show evidence for linkages between genetic variations in hyperoxia tolerance, H_2_O_2_ signaling, and pyrenoid morphologies.

## INTRODUCTION

The maximal primary productivity of algae is often determined by the efficiency of photosynthesis, which is strongly impacted by environmental factors. In turn, the products of photosynthesis can also impact local environmental conditions, leading to feedback- (or self-) limitations (Livansky, 1996; Pulz, 2001; Raso et al., 2012; Torzillo et al., 1998; Vonshak et al., 1996; Weissman et al., 1988). One important, but relatively little studied feedback factor is hyperoxia, which results when O_2_ is emitted as a by-product of photosynthesis more rapidly than it diffuses away or is consumed by respiration. Microalgae cultures are often observed with dissolved oxygen levels of up to 100-400% of air – or even higher (Peng et al., 2013), especially when the local supply of inorganic carbon (C_i_) is high but consumption or diffusion of O_2_ slow. In some species, hyperoxia constitutes a major hurdle in achieving low cost, highly productive micro-algae farms (Peng et al., 2013). Hyperoxia has been directly associated with loss of productivity in a wide range of algal and cyanobacterial species, including *Nannochroposis* (Raso et al., 2012), *Chlamydomonas reinhardtii* (Kliphuis et al., 2011), *Neochloris oleabundans* (Peng et al., 2016; Sousa et al., 2012), *Chlorella sorokiniana* (Ugwu et al., 2007) and *Spirulina* (Vonshak et al., 1996).

Despite being recognized as a problem, how hyperoxia interferes with photosynthetic growth is not fully understood, and various mechanisms have been proposed, including reactive oxygen (ROS)-induced damage to the photosynthetic machinery, membrane structure, DNA, and other cellular components (Marquez et al., 1995; Santabarbara et al., 2002; Ugwu et al., 2007). Another mechanism by which high O_2_ has been proposed to decrease productivity is photorespiration, a process initiated when the ribulose bisphosphate carbolylase/oxygenase (rubisco) enzyme fixes O_2_ rather than CO_2_, resulting in the production of the toxic side product phosphoglycolate, which is detoxified through the photorespiratory pathway, at the cost of lost energy and the release of fixed carbon (Bauwe et al., 2010). The intermediates of photorespiration can also form rubisco inhibitors (Kim and Portis, 2004) that can further slow photosynthesis. If rubisco becomes inactivated, then ROS accumulation can lead to chlorosis and cell death, particularly in high light (Spreitzer and Mets, 1981).

Because photorespiration depends on competition between CO_2_ and O_2_ at rubisco, it can also contribute to loss of productivity under low inorganic carbon (Wang et al., 2015). The current atmospheric CO_2_ concentration is well below the saturated concentration for rubisco’s carboxylase activity (Raven et al., 2008) and CO_2_ diffuses through aquatic environments 10,000 times shower than in air (Moroney and Ynalvez, 2007). Thus many aquatic phototrophs, including the chlorophyte *Chlamydomonas reinhardtii*, possess carbon concentrating mechanisms (CCMs) that concentrate CO_2_ above its K_M_ at rubisco, which has been reported to be 29 μM (Jordan and Ogren, 1981) to 57 μM (Berry et al., 1976), in order to increase the relative rates of carboxylation relative to oxygenation (Aizawa and Miyachi, 1986; Badger et al., 1980).

The expression and function of green algal CCMs in eukaryotic algae is highly regulated; cells grown on or below air levels of CO_2_ (0.04%) develop active CCMs (Aizawa and Miyachi, 1986; Badger et al., 1980), whereas those grown with high CO_2_ levels lack them, and thus show low apparent affinities for CO_2_. Cells grown at high CO_2_ and rapidly transferred to low CO_2_ show strong inhibition of photosynthesis (Badger et al., 1980; Spalding et al., 1983) until the CCM is induced and activated (Aizawa and Miyachi, 1986; Badger et al., 1980; Manuel and Moroney, 1988). The CCM in Chlorophytes involves a large number of components, including proteins that serve enzymatic and structural functions as well as a starch sheath that surrounds the pyrenoid, forming a subcellular compartment which acts as a trap to concentrate pumped inorganic carbon near localized rubisco (Mackinder et al., 2017; Ramazanov et al., 1994; Wang et al., 2015). The pyrenoids are penetrated by tubule-like extensions of the thylakoid membranes, which are thought to supply CO_2_ to the trapped rubisco by dehydration of luminal HCO_3_^-^ (Engel et al., 2015; Mitra et al., 2004; Moroney and Ynalvez, 2007).

Extensive genetic and biochemical studies have identified a large number of other components essential for CCM function (Goodenough and Levine, 1970; Henk et al., 1995; Itakura et al., 2019; Spalding et al., 1983; Toyokawa et al., 2020). Of particular interest to the current work are factors that contribute to the pyrenoid compartment itself, especially those that affect the localization of rubisco within its starch sheath or those that modify the structure of the starch sheath. A range of mutants in diverse genetic components fail to form pyrenoids (Goodenough and Levine, 1970; Henk et al., 1995; Spreitzer et al., 1985), or have altered pyrenoid ultrastructure with disorganized or missing starch sheaths (Henk et al., 1995; Itakura et al., 2019; Toyokawa et al., 2020). These mutants tend to require high CO_2_ for growth, emphasizing the importance of the pyrenoid structure for the function of the CCM. However, the pyrenoid is not necessary in all cases for survival under low CO_2_, as some species of *Chloromonas*, despite lacking pyrenoids, have functioning CCMs (Morita et al., 1998).

In this work, we explore the importance of an aspect of the CCM, in particular the pyrenoid, in responses to hyperoxia, rather than low Ci availability. Since both low CO_2_ and hyperoxia involve a lowering of the CO_2_:O_2_ ratio, we hypothesized that 1) the pyrenoid is induced by hyperoxia; 2) that differences in its induction and/or formation can be related to hyperoxia tolerance. Furthermore, since both low CO_2_ and hyperoxia result in increased photorespiration, we hypothesized that 3) the signal for pyrenoid formation might be a by-product of photorespiration, H_2_O_2_. In order to address these hypotheses, we examined two natural isolates of *Chlamydomonas* with varying tolerances to hyperoxia, and their progeny, with the goal of better understanding the physiological mechanisms that underly responses to hyperoxia, and potentially the genetic loci that underlie such mechanisms. Understanding such traits can give insights into the mechanisms and tradeoffs of adaptations for specific environmental niches. By extension, such traits and tradeoffs have strong relevance to applications ranging from algae cultivation to bioengineering crops for increase productivity (Long et al., 2015).

## METHODS

### Chlamydomonas Strains and Mating

Strain CC-2343 (mt+), in a search for strains resistant to heavy metals CdCl_2_ and HgCl_2_, was isolated from soil in Melbourne, Florida in 1988 (Spanier et al., 1992). Strain CC-1009 (mt-) is a wild type strain tracing back to the 1945 collection by G.M. Smith, collected from Amherst MA, but has been a separate line from the c137c (CC-124 and CC-125) and Sagar (CC-1690) since about 1950. CC-5357 was generated by Luke MacKinder in the laboratory of Martin Jonikas (Mackinder et al., 2016). These (Supplementary Material Figure 1) and other strains were obtained from the Chlamydomonas Resource Center (https://www.chlamycollection.org). CC-2343 and CC-1009 were mated using an established protocol (Jiang and Stern, 2009).

### Growth and Biomass

Cultures (i.e. CC-2343, CC-1009, and progeny) were grown autotrophically in environmental photobioreactors (ePBRs) (Lucker et al., 2014), or in some cases in 125 mL Erlenmeyer flasks, in either a medium called 2NBH (Davey et al., 2012), which is a modified Bristol’s medium (Supplementary Material Table 1), or (i.e. CC-5357 and other strains descendant from CC-4533) in HS (Sueoka’s high salt) medium (Sueoka, 1960) because of their requirement for ammonia rather than nitrate as a nitrogen source. When grown in ePBRs, culture density was maintained by turbidostat-controlled automatic dilution, adjusted to give chlorophyll concentrations of approximately 3 μg/mL. The media filled the columns to 15 cm in height, bringing the total volume to 330 mL. Following inoculation, all cultures were maintained at least three days at constant chlorophyll prior the measuring of productivity. Standard illumination was provided on a 14:10 hour (light:dark) sinusoidal diurnal cycle, with the peak light intensity of 2000 μmol m^−2^ s^−1^. Gas was filtered with using a HEPA-Cap disposable air filtration capsule (Whatman^®^, #67023600), and bubbled through a 5 mm gas dispersion stone with a porosity of 10-20 microns at a flow rate of 350 ml/min.

In our ePBRs, we used a series of sparging protocols to establish a range of CO_2_ and O_2_ levels as well as to simulate fluctuations in CO_2_ that might occur during production culturing, including: 1) rapid sparges (one min on and one min off) during illumination; 2) “raceway sparges”, one min sparge each hour during illumination. For normoxic conditions, sparge gas was 5% CO_2_, 21% O_2_, balance N_2_. For “hyperoxia” treatments, the sparge gas was 5% CO_2_ and 95% O_2_.

Biomass productivity in units of g·m^2^·day^−1^ was estimated by multiplying the volume of eluted culture as a result of turbidostatic dilutions by the measured Ash Free Dry Weight (AFDW) per unit volume, then normalizing to m^2^ by dividing by the surface area of the ePBR water column. The column height was 15 cm and the surface area is 26.6 cm^2^. Unless noted otherwise, all experiments were done in biological triplicate, each separate bioreactor or flask representing a different biological replicate.

When grown in the Erlenmeyer flasks, the cultures were grown in batch mode (Anderson, 2005) under ~80 μmoles m^−2^ s^−1^ of light and bubbled continuously via a glass Pasteur pipette with 5% CO_2_.

In our aerophilic, mixotrophic assays (i.e. Figure 4), cells were grown at steady state in 2NBH media in photobioreactors and then counted using a Beckman Coulter Z2 Coulter Counter at sizes between 3-10 microns. 50000, 5000, 500, 50 cells were then spotted onto Tris Acetate Phosphate (TAP) plates (Gorman and Levine, 1965) and grown under 80 μmoles m^−2^ s^−1^ of light.

### Estimation of culture bicarbonate concentrations

Dissolved bicarbonate levels were estimated using an approach based on the release of CO_2_ upon acidification of the media (Hawkes et al., 1993) using an in-house built instrument consisting of a 250 mL sealed glass reactor (a standard canning jar, Mason, USA) that houses a small but sensitive atmospheric CO_2_ sensor (S8, www.senseair.com), a 3 cm long Teflon-coated magnetic stir bar and a small septum for introducing reagents. During experiments, the output of the CO_2_ sensor was continuously collected at a rate of 1Hz using a microcontroller (Teensy 3.2, PJCR, Sherwood, OR, USA) and analyzed with a Python Jupyter (www.jupyter.org) notebook. The experiments started with the addition of 10 mL of sample and continuous stirring (approximately 30 Hz rotation frequency). Similar results were obtained when samples were drawn directly from the ePBR or passed through a 4 micron filter to remove cells. After introducing the sample, the system was allowed to equilibrate for 3 min., at which time the sample was acidified to pH < 4 by addition of 200 μL of 1N HCl. The acidification leads to hydrolysis of HCO_3-_ to CO_2_ + H_2_O, resulting in release (outgassing) of CO_2_ into the chamber. To account for differential partitioning of CO_2_ into the medium and atmosphere, responses were then calibrated by spiking the samples with a known concentrations of sodium bicarbonate.

### Microscopy

At each time point of interest, 1 mL of culture (at ~3 μg/mL chlorophyll) were removed from our bioreactors, placed in Eppendorf tubes and mixed with 2 μ**L** of Lugol’s Solution (Sigma-Aldrich, cat. no. L6146) before being viewed in a Leica DMi90 inverted light microscope.

Transmission Electron Microscopy (TEM) was performed using a JEOL 1400 Flash instrument, and images were photographed with a Metattaki Flash cMOS camera. To prepare cells for microscopy, samples were resuspended in 2.5% glyceraldehyde in cultures of 2NBH media, and then treated as previously described (Du et al., 2018). Morphometric analysis of the images was conducted using a previously described protocol (Weibel, 1969). Briefly, a grid was placed over the cells, to scale, and then the number of intersecting points within a particular morphology characteristic were quantified.

Subcellular localization of Rubisco labeled with Venus fluorescence protein was imaged using an Olympus FluoView 1000 Confocal Laser Scanning Microscope, configured on an Olympus IX81 inverted microscope using either a 60x PlanApo (NA 1.42) oil objective or a 100x UplanApo (NA 1.40) oil objective. Venus fluorescence protein was excited using the 515 nm Argon laser emission line, and fluorescence emission was detected using a 530-620nm band pass filter. We also repeated the analysis using a Nikon A1 Confocal Laser Scanning Microscope, configured on a Nikon Ti Eclipse inverted microscope using a 100x Apo TIRF (NA 1.49) oil objective. Venus fluorescence protein was excited using the 515 nm Argon laser emission line, and fluorescence emission was detected using a 530-600nm band pass filter. Transmitted laser light was simultaneously collected using brightfield optics. Confocal Z-series through the thickness of the algal cells were collected in 0.5 micron increments, typically through a 5 micron thickness, and the Z-stacked images were compressed into a 2D image, displayed as a Maximum Intensity Projection.

Confocal work to probe cellular reactive oxygen species (ROS) production was performed on the Olympus confocal microscope setup described above, using methods previously described (Du et al., 2018).

### H_2_O_2_ measurements

For H_2_O_2_ measurements, cells were treated with reagents of an Amplex Red Hydrogen Peroxide/Peroxidase Assay Kit (Molecular Probes/Invitrogen, Carlsbad, CA, USA), as has been used by previous researchers (Lin et al., 2013). In brief, 5 mL of the culture was collected by centrifugation, and the pellet was flash frozen in liquid nitrogen. The cells were then broken in 1 mL of 1 x reaction buffer from the assay kit, ground with glass beads, and briefly sonicated. The mixture was then centrifuged and the supernatant was then used to measure the cellular H_2_O_2_ concentrations after incubation with horseradish peroxidase at 25 °C for 30 minutes. The H_2_O_2_ concentrations were determined by a standard curve developed using 0.25 – 2.5 μM and normalized by calculating the amount of protein in the extract using a standard Bradford Assay (Bradford, 1976) with Bradford Reagent (Sigma-Aldrich, cat no. B6916).

### Immonoblotting

Total protein was extracted with 2X Laemmli buffer (Sigma-Aldrich, cat no. 33401) containing 5% β-mercaptoethanol by boiling at 95°C for 5 minutes. Protein concentrations were determined using the RC DC Protein Assay kit (Bio-Rad, cat. no. 5000122). Quality of protein samples were verified by loading equal protein amounts into an SDS-PAGE gel before capillary immunoblotting. Protein quantities were measured following the manufacturer’s instructions using the Wes (Protein Simple, www.proteinsimple.com) on the 12-230 kD Master Kit (Protein Simple) with Agrisera and PhytoAB antibodies (see Supplementary Table 3 for antibodies used).

### Rubisco Activity Assay

Rubisco enzymatic activity was assayed using an established protocol (Li et al., 2019; Roeske and Oleary, 1985; Sharkey et al., 1986), with slight adjustments to make the protocol suitable specifically for *Chlamydomonas*. Briefly, cultures were harvested from the bioreactors, flash frozen in liquid nitrogen, and stored at −80°C. Just prior to assay, samples were suspended in extraction buffer [50 mM 4-2(2-hydroxyethyl)-1-piperazine propane sulfonic acid (EPPS), pH 8, 30 mM NaCl, 10 mM mannitol, 5mM MgCl_2_, 2mM EDTA, 5 mM DTT, 0.5% (v/v) Triton X-100, 1% polyvinylpolypyrrolidone (PVPP), 0.5% casein, and 1% protease inhibitor cocktail (P9599; Sigma-Aldrich)], sonicated and vortexed to extract proteins. Aliquots of 20 μL of the extract was added to 80 μL of assay buffer [50 mM 4-2(2-hydroxyethyl)-1-piperazine propane sulfonic acid, pH 8, 5 mM MgCl_2_, 0.2 mM EDTA, 0.5 mM Ribulose bisphosphate, and 15 mM NaH^12^CO_3_, and 0.3 mM H^14^CO_3_^-^]. The suspensions were vortexed for three seconds, incubated for one minute, and then the reaction was halted by adding 100 μL of 1 M formic acid. The resulting acidification liberates unfixed inorganic C by converting HCO_3-_ to CO_2_ which escapes from the buffer. The mixtures were vortexed again for 3 seconds and then dried on a hotplate at 75°C. For measurements of total rubisco activity, the extracts were pre-incubated with activation solution (to give final concentrations of 20 mM MgCl_2_, 15 mM H^12^CO_3_, and 61 mM 6-phosphogluconate) for ten minutes before being mixed with the assay buffer. The amount of fixed radioactivity was determined using a liquid scintillation counter (TriCarb^®^ 2800TR, Perkin Elmer). Each day, radioactivity in 10 μL of the assay buffer was counted to determine specific activity. Based on 1mCi = 2.22 x 10^9^ disintegrations min^−1^, initial and total rubisco activity was calculated as expressed as μmol m^−2^ s^−1^. The rates were divided by 0.943 to account for the discrimination against ^14^C (Li et al., 2019; Roeske and Oleary, 1985). Three biological replicates were run per treatment or condition, and each biological replicate constituted three technical replicates.

### Oxygen Evolution and Φ_II_

Cell suspensions growing at steady state were removed from the bioreactors and concentrated in fresh media to 50 μg/mL of chlorophyll in a cuvette. To drive out the oxygen, the cultures were then sparged with 1% CO_2_ and 99% N_2_ gas. Subsequently also supplemented with 6.25 mM sodium bicarbonate, the cultures were illuminated with approximately 750 μmol photons m^−2^ s^−1^ of photosynthetically active radiation (PAR), measured using a submersible spherical micro quantum sensor (US-SQS/L, Walz) attached to a light meter (Li-250A, LiCor) from two red LEDs (emission at approximately 630 nm) aimed at opposing sides of the cuvette. Oxygen evolution was then measured using a Neofox oxygen sensor via a fiber optic fluorescent probe by Ocean Optics (Dunedin, Florida). The **Φ_II_** measurements of the TAP plates were made in in our dynamic environmental photosynthesis imager (DEPI), using methods described previously in detail (Cruz et al., 2016) but, to avoid direct reflection of the measuring light into the camera, the plates were tilted by approximately 5° from the horizontal position.

## RESULTS

### Hyperoxia differentially affects rubisco activity in the tolerant and sensitive lines

In an initial screen of sequenced *Chlamydomonas* isolates (Jang and Ehrenreich, 2012), we found two with contrasting tolerances to hyperoxia, with strain CC-1009 relatively tolerant to hyperoxia, continuing to grow, albeit at a suppressed rate, when exposed to 95% oxygen and 5% CO_2_, while CC-2343 showed severely suppressed growth and eventual chlorosis or photobleaching in our ePBRs (Hall, 2017). Qualitatively, this varying tolerance was also observed when cultures were continuously sparged in batch culture (Supplementary Material Figure 2), when the cultures were CO_2_ saturated, indicating that the differential sensitivity was caused by hyperoxia rather than depletion of inorganic carbon sources (see also results on rapid sparging in the ePBR system, below).

Spreitzer and Mets (1981) found that rubisco activity-deficient mutants exhibited chlorotic phenotypes similar to those observed with CC-2343 under hyperoxia, so we measured rubisco activity of both strains prior to and after 31 hours exposure to hyperoxia (Supplementary Material Figure 3). Rubisco activity was measured immediately after isolation to estimate steady-state activity at the time point of interest, which is controlled by both the total enzyme content and the fraction of the enzyme in the inactive state related to carbamylation state or the presence of inhibitors (Li et al., 2019b; Roeske and Oleary, 1985). Pre-incubating for ten minutes in the presence of MgCl_2_, HCO_3_^-^, and 6-phosphogluconate (6-PG) promotes activation of the enzyme by stabilizing the Enzyme-CO_2_-Mg-Complex of rubisco, allowing for the estimation of the maximal rubisco activity (Badger and Lorimer, 1981; Chu and Bassham, 1973; Matsumura et al., 2012). Using this method, we estimate that, under atmospheric levels of O_2_, approximately 60% of the enzyme was in its active form for both CC-1009 and CC-2343. After 31 hours of exposure to hyperoxia, the total (maximal) activity of rubisco decreased in both lines, by about 10% and 28% in CC-1009 and CC-2343, respectively. However, in the case of CC-1009, the loss in total activity was compensated for by a large increase (to about 95%) in the fraction of active enzyme, leading to an overall increase of about 23% in steady-state activity. By contrast, the fraction of activated rubisco was unchanged in CC-2343, leading to an overall decrease of about 13% in steady-state activity.

Although we cannot ascribe the differences in photosynthetic phenotypes solely to rubisco deactivation, these results do suggest that the CO_2_/O_2_ concentrations or metabolic environments near rubisco are different under hyperoxia in the two lines. Apart from the metabolic environment, the activation state of rubisco can also be affected by the levels or activity of the rubisco activase (Pollock et al., 2003). But we found no consistent differences in the cellular contents of the rubisco activase protein between the cell lines (Supplementary Material Figure 4). Another important factor that could affect the activity of rubisco and its metabolic environment in *Chlamydomonas* is its pyrenoid, a distinct, well-structured starch sheath surrounding localized rubisco that, under low CO_2_, plays a key role in trapping CO_2_ in the CCM (del Campo et al., 1995; Harris, 1989; Harris, 2009; Ramazanov et al., 1994). It has also been proposed to shield rubisco from high O_2_ levels generated by PSII (McKay and Gibbs, 1991; Toyokawa et al., 2020). We thus initially hypothesized that: 1) the pyrenoid could be important for responses to hyperoxia and 2) differences in pyrenoid structure or regulation may then contribute to the distinct photosynthetic responses in the two lines. Consistent with these hypotheses, under saturating CO_2_ conditions (Supplementary Material Figure 5), the pyrenoid starch sheaths are not clearly discernable, in agreement with previous research showing that the pyrenoid starch sheath is not expressed under Ci replete conditions (Borkhsenious et al., 1998; Ramazanov et al., 1994). Exposure to hyperoxia (95% O_2_ and 5% CO_2_), both when sparged rapidly (a square wave cycle one minute sparge and one minute rest, see Materials and Methods) (Supplementary Material Figure 6 & 7) or under the raceway sparging regime (one minute sparge every hour, see Figure 1) strongly induced starch sheath formation in our strains, but with genotype-dependent morphologies (Figure 1). The tolerant line, CC-1009, exhibited clearly defined, continuous starch sheath rings around its pyrenoid compartment, punctuated only in places where thylakoid tubules enter the pyrenoid matrix. By contrast, CC-2343 showed a more fragmented and porous structure, with gaps in the (thin section, TEM) images that were not clearly association with tubules.

**Figure 1:**
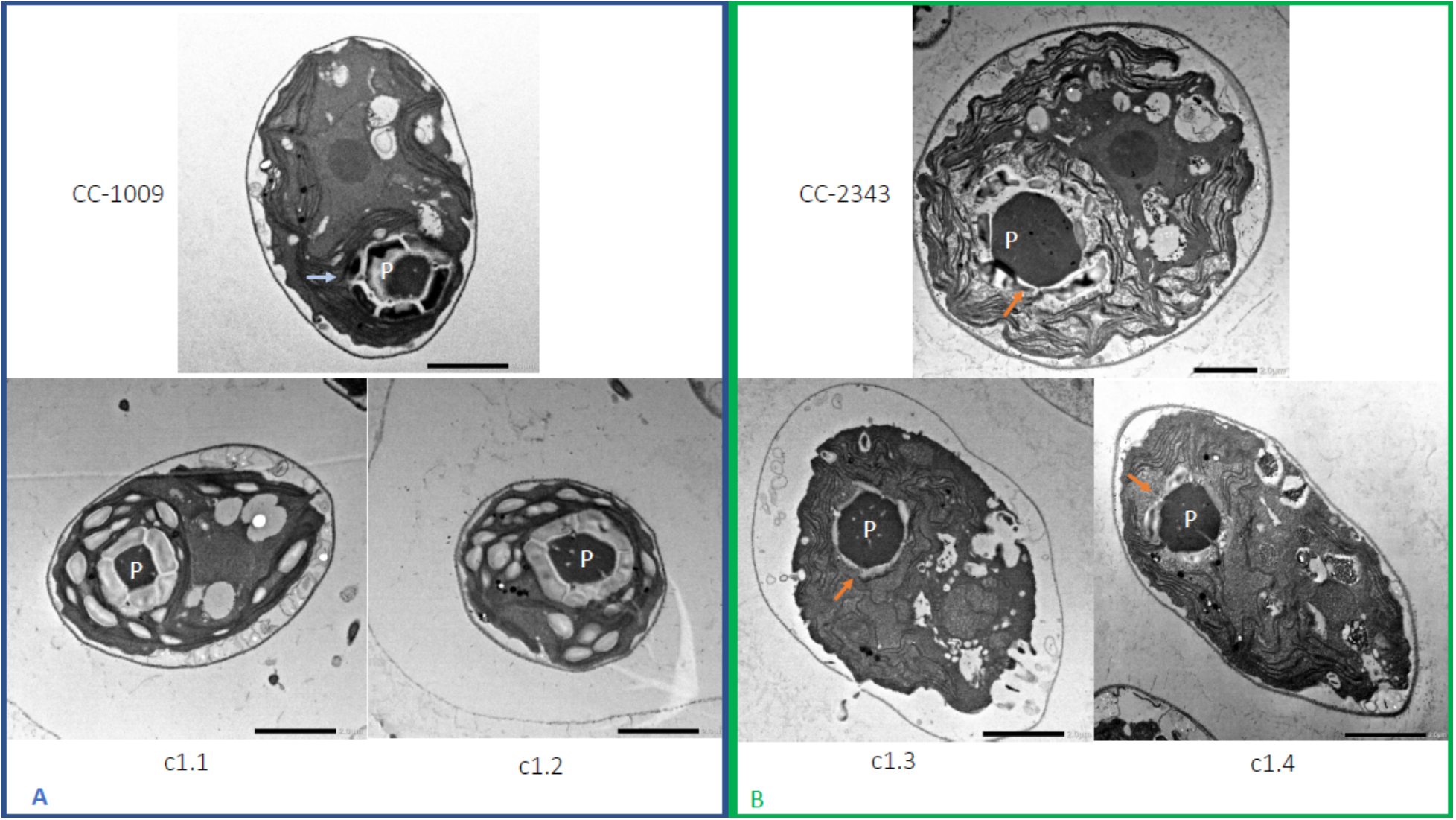
Representative TEM images of *Chlamydomonas* strains, the parents (CC-1009 & CC-2343), as well as their progeny c1_1, c1_2, c1_3, c1_4 after being exposed to hyperoxia for six hours in the light. Clear differences were observed between the pyrenoids of the cells in Panel A and Panel B, with the cells in Panel A showing more robust, continuous, sealed pyrenoids. Prior to switching the gas to hyperoxia (i.e. 95% O_2_ and 5% CO_2_) cells had been grown in steady state conditions, with 5% CO_2_ with 14:10 hour (light:dark) sinusoidal illumination with peak light intensity of 2000 μmoles m^−2^ s^−1^, in minimal 2NBH media. Cells here were fixed at 11:00 am, at 1945 μmoles m^−2^ s^−1^. Pyrenoids are labeled with “P.” Orange arrows point out gaps in starch sheaths. Scale bar = 2 μm.

To test if decreased CO_2_ or inorganic carbon levels could account for induction of pyrenoid synthesis, we directly assayed levels at various times during the sparge cycle, using the method described in Materials and Methods. For the rapid sparging protocol, the estimated [HCO_3-_] under both normoxia and hyperoxia remained between 2-3 mM, and for the “raceway” sparging protocol, between 1.3-1.7 mM a few minutes after sparging and 1.0-1.4 mM just prior to the following sparge. In all cases, the pH of the medium remained below 7.4, and thus based on the known pK_a_ values for the CO_2_/bicarbonate system, we estimate the lowest CO_2_ levels experienced by the cultures, which occurred under raceway sparging conditions, remained above 100 μM, above the K_m_ of rubisco (~29 μM – 57 μM) in *Chlamydomonas* (Berry et al., 1976; Jordan and Ogren, 1981) and in excess of the concentration found by Toyokawa et al. (2020) to induce the formation of the pyrenoid starch sheath (2.1-3.1 μM). The CCM in *C. reinhardtii* is typically induced when the concentration of CO_2_ in the air bubbled through the culture is decreased to around 0.5% or lower (Vance and Spalding, 2005).

Similar genotype-dependent pyrenoid morphologies were also observed, in a 2:2 segregation pattern, in four daughter cells dissected from a single tetrad (Figures 1). Two of the progenies, designated c1_1 and c1_2, when exposed to hyperoxia, developed completely sealed and robust rings, like CC-1009, while two others, designated c1_3 and c1_4, showed fragmented, porous structures, like CC-2343. These differences were even more apparent after 31 hours of hyperoxia (Figure 2). Strains with fragmented pyrenoids (CC-2343, c1_3 ad c1_4) showed an abrupt inhibition of growth after one day of exposure to hyperoxia, whereas those with sealed pyrenoids (CC-1009, c1_1 and c1_2) continued to grow rapidly and produce biomass (Figure 3). Both progeny with fragmented pyrenoid sheaths grew even more slowly than the sensitive parent, CC-2343. On the other hand, those progeny with sealed pyrenoids (c1_1 and c1_2) initially grew more slowly than CC-1009, but maintained steady growth even on the fourth day of hyperoxia (Figure 3). These results suggest that the ultrastructural differences in the pyrenoid starch sheath (Figure 1 and Figure 2) are related to the observed tolerances of growth to hyperoxia in both parent and progeny lines (Figure 3). However, the differences among the tolerant and sensitive lines, particularly the observation that the progeny have phenotypes more extreme than those of the parent lines, imply that additional genetic factors (beyond those that control pyrenoid morphology) likely contribute to productivity under hyperoxia.

**Figure 2:**
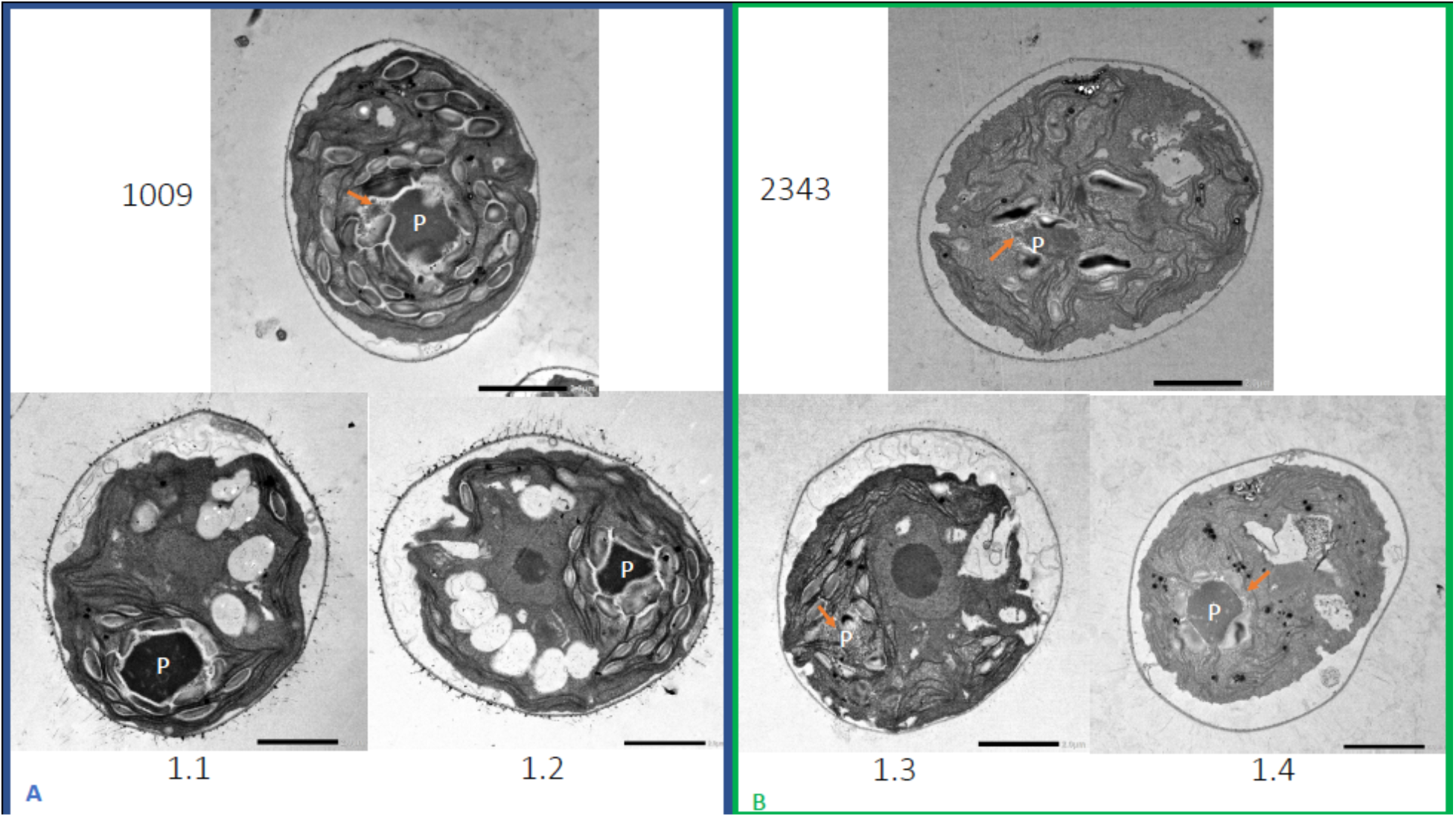
Representative TEM images of *Chlamydomonas* strains, of the parents (CC-1009 & CC-2343), as well as their progeny c1_1, c1_2, c1_3, c1_4 after being exposed to hyperoxia for 31 hours in the light. None of the dozens of cells we observed in CC-2343, c1_3, or c1_4 (Panel B) exhibited a continuous, sealed starch sheath around the pyrenoid as often exhibited by CC-1009, c1_1, and c1_2 (Panel A). Prior to switching the gas to hyperoxia (i.e. 95% O_2_ and 5% CO_2_) cells had been grown in steady state conditions, with 5% CO_2_ with 14:10 hour (light:dark) sinusoidal illumination with peak light intensity of 2000 μmoles m^−2^ s^−1^, in minimal 2NBH media. Cells here were fixed at 11:00 am, at 1945 μmoles m^−2^ s^−1^. Pyrenoids are labeled with “P.” Orange arrows point out gaps in starch sheaths. Scale bar = 2 μm.

**Figure 3:**
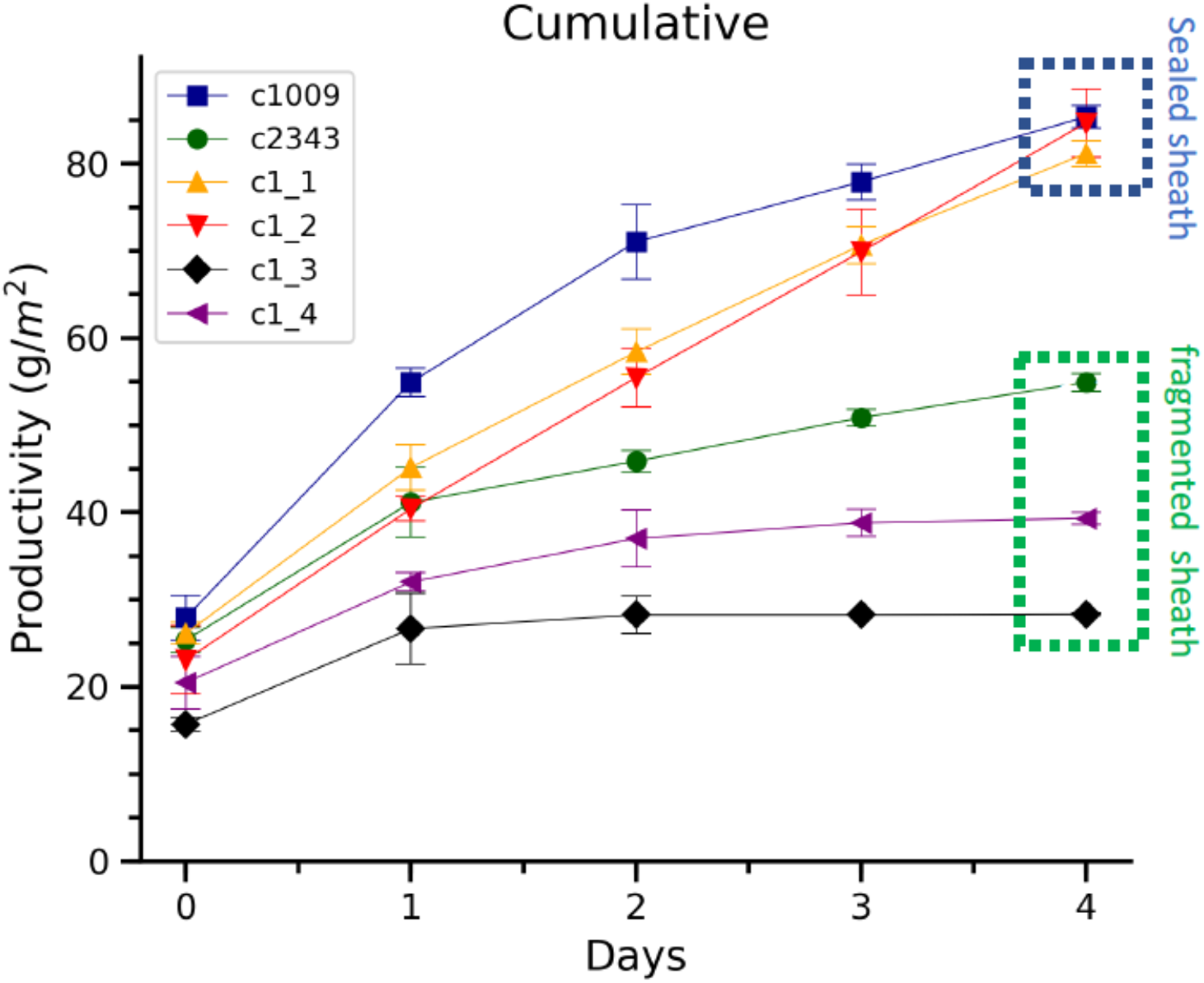
Cumulative biomass productivity following switching the bioreactors over to hyperoxia at dawn on day 0. Strains CC-1009, c1_1, and c1_2, which all showed continuous, sealed pyrenoids at 6 hours (see Figure 1, Panel A) all continued to accumulate biomass after three days of hyperoxia, while CC-2343, c1_3, and c1_4, which had fragmented, porous pyrenoid (Figure 1, Panel B) structures, did not, with daily productivities hovering at zero. Visual inspection (via light microscopy) also revealed that the cultures of the intolerant lines by day 3 consisted of severely stressed or dead cells, while the tolerant lines showed cells with continued viability. Prior to exposure to hyperoxia, cultures were grown at steady state (with 5% CO_2_ with 14:10 hour (light:dark) sinusoidal illumination with peak light intensity of 2000 μmoles m^−2^ s^−1^, in minimal 2NBH media) and at Day 0, the gas was switched to hyperoxia not at midnight but at dawn. Even at steady state, c1_3 had lower growth than the other strains, although this was not true when grown at other conditions (i.e. see Figure 4). Error bars represent standard deviation for 3 separate reactor experiments. By day 3, c1_3 always ceased growth. Even though productivities had just begun to decline at 6 hours, the pyrenoid structure (i.e. sealed vs. porous Figure 1) paralleled the eventual tolerances.

We also plated cultures of the CC-2343 and CC-1009 and the four progenies on TAP agar plates. Interestingly, growing the cells under aerophilic, mixotrophic conditions, we found that CC-2343 and the progeny that were intolerant to hyperoxia (c1_3 and c1_4) grew more rapidly than the hyperoxia tolerant lines which had exhibited the sealed, continuous pyrenoid starch sheaths (CC-1009, c1_1, c1_2) (Figure 4), despite exhibiting similar Φ_II_ values (Supplementary Material Figure 8). These observations support that the growth advantage on the TAP plate was related to carboxylation, at least more likely than an aspect of the light reactions. In addition, when we grew the parent cells under steady state supplemented with 5% CO_2_, CC-2343 synthesized more starch (Supplementary Material Figure 9).

**Figure 4:**
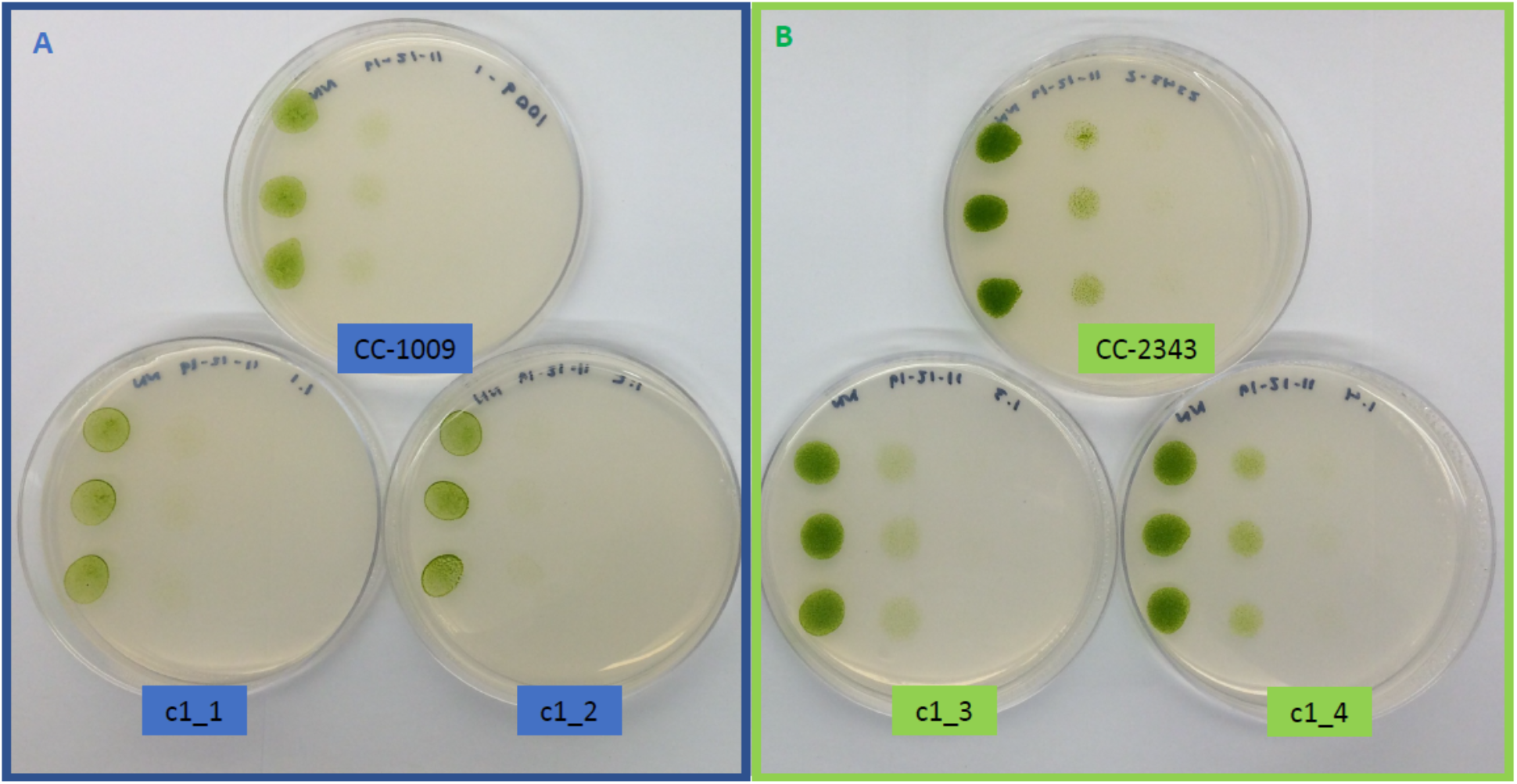
Growth of parents (CC-1009 and CC-2343) and F1 tetrad offspring following the serial dilution, by column, of a cultures on TAP agar plates. Rows are replicate dilutions. Photo was taken 3 days after plating. The oxygen intolerant lines (Panel B) all clearly grew better than the tolerant line (Panel A). See Figure 3 for graph of oxygen tolerance.

Consistent with studies which have shown the pyrenoid is light dependent (Kuchitsu et al., 1988; Lin and Carpenter, 1997), CC-1009, CC-2343, and the F1 tetrad offspring also lost visible pyrenoid structures after dark exposure during the night (sparging once every hour with 5% CO_2_ in air), and the pyrenoid starch sheaths did not appear fully formed during the morning when PAR was low (Figure 5). As the light levels increased over 6 hours, though, the pyrenoid structures still formed under raceway sparging. Rather than a specific light level, this could be because it takes several hours to form pyrenoid structures and that photosynthesis is likely required. Under these conditions, CC-1009, c1_1, and c1_2 exhibited more tightly structured pyrenoids (Figure 6), though the differences were not as great as those exhibited under hyperoxia (Figure 1).

**Figure 5:**
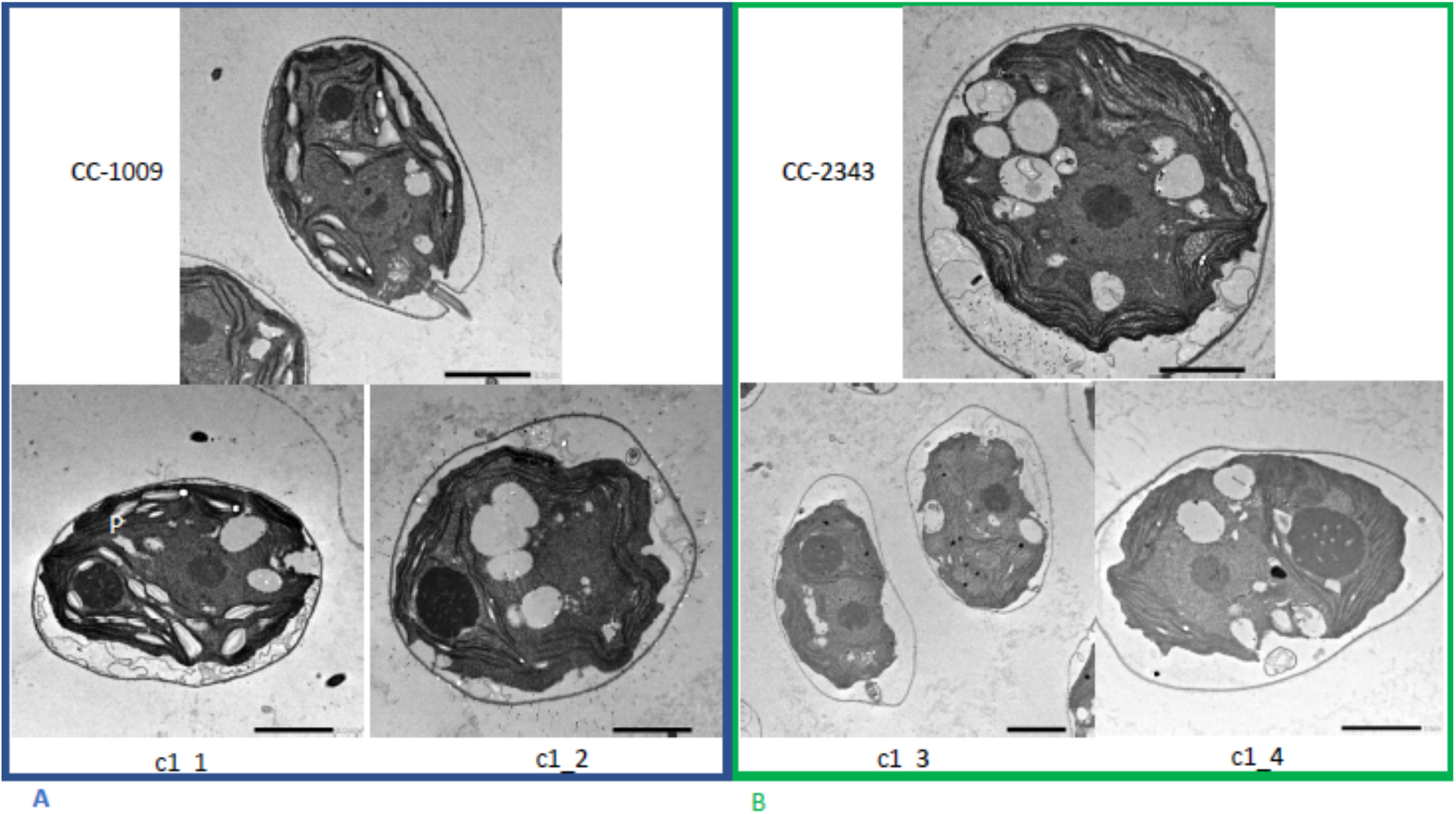
Representative TEM images of *Chlamydomonas* strains, the parents (CC-1009 & CC-2343), as well as their progeny c1_1, c1_2, c1_3, c1_4, having been sparged with CO_2_ for 30 seconds every hour during the night. Cells had been grown in steady state conditions, with 5% CO_2_ with 14:10 hour (light:dark) sinusoidal illumination with peak light intensity of 2000 μmoles m^−2^ s^−1^, in minimal 2NBH media. Cells here were fixed two hours after dawn, at 7:00 am, when light levels were 825 μmoles m^−2^ s^−1^. Pyrenoids appeared absent or mostly unsheathed in these conditions. Scale bar = 2 μm.

**Figure 6:**
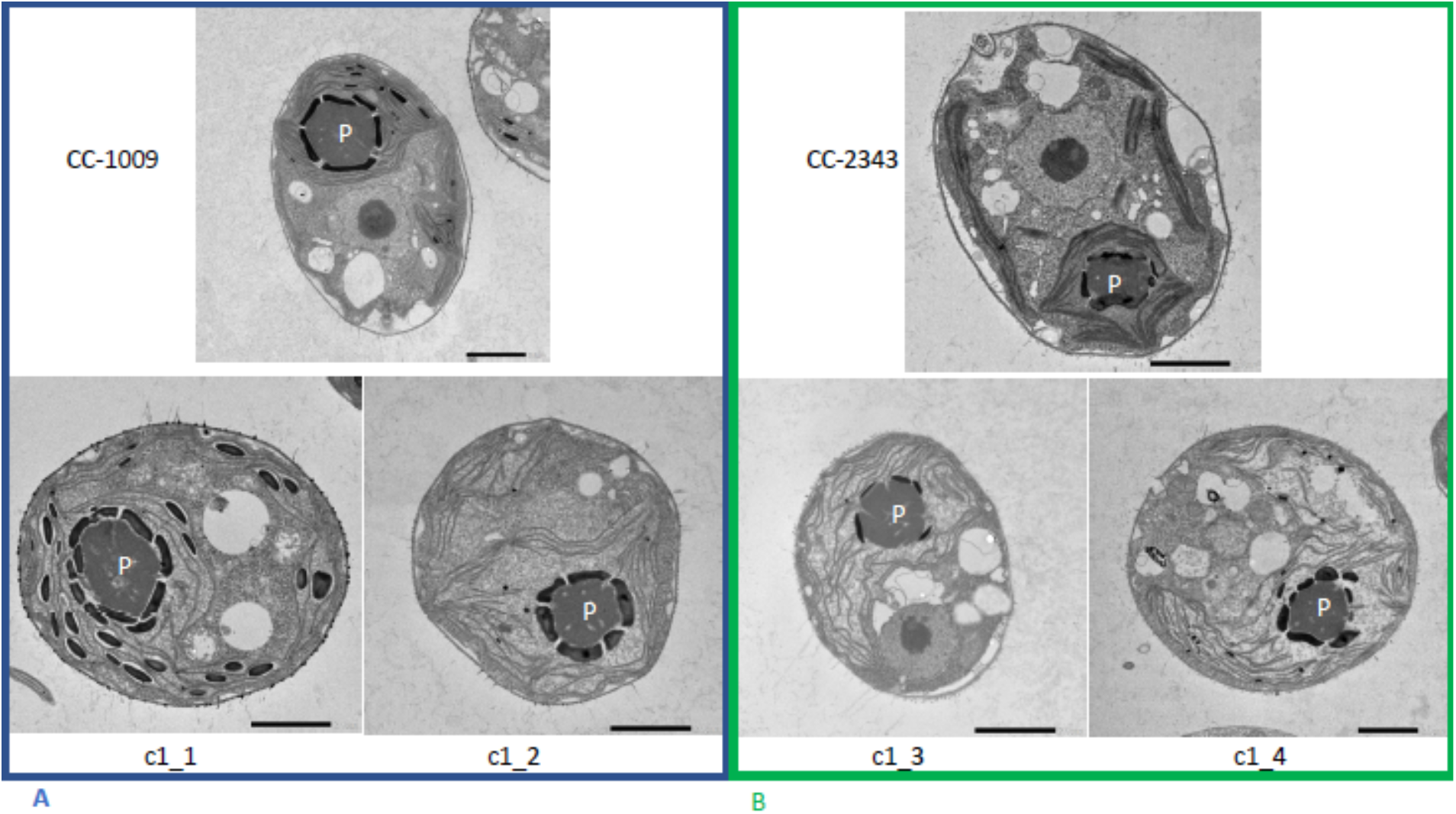
Representative images of *Chlamydomonas* strains, the parents (CC-1009 & CC-2343) as well as their progeny c1_1, c1_2, c1_3, and c1_4. Images taken while cells were being grown at steady state, being sparged with 5% CO_2_, 95% O_2_, with 14:10 hour (light:dark) sinusoidal illumination with peak light intensity of 2000 μmoles m^−2^ s^−1^, in minimal 2NBH media. Differences were observed between the pyrenoids of the cells in Panel A and Panel B. Cells here were fixed at 11:00 am, at 1945 μmoles m^−2^ s^−1^. Pyrenoids are labeled with “P.” Scale bar = 2 μm.

Consistent with previous work (Borkhsenious et al., 1998), pyrenoid formation was observed in all lines when cells were grown at high light and low CO_2_, but with some differences in morphology among the lines. After exposure to low CO_2_ (i.e. ambient air) for 6 hours, c1_1, c1_2 showed tightly closed sheath morphology similar to CC-1009 (Figure 7, Panel A). However, after 31 hours of exposure to low CO_2_, the genotype differences in morphology became less apparent as all lines made starch sheaths of some integrity (Figure 8). All lines also grew similarly under ambient CO_2_ in flasks under approximately 85 μmol photons m^−2^ s^−1^ (Supplementary Material Figure 10). Taken together, these results suggest that low inorganic carbon, high O_2_ and high light can all promote synthesis of the starch sheath, and that genetic variations modulate these responses.

**Figure 7:**
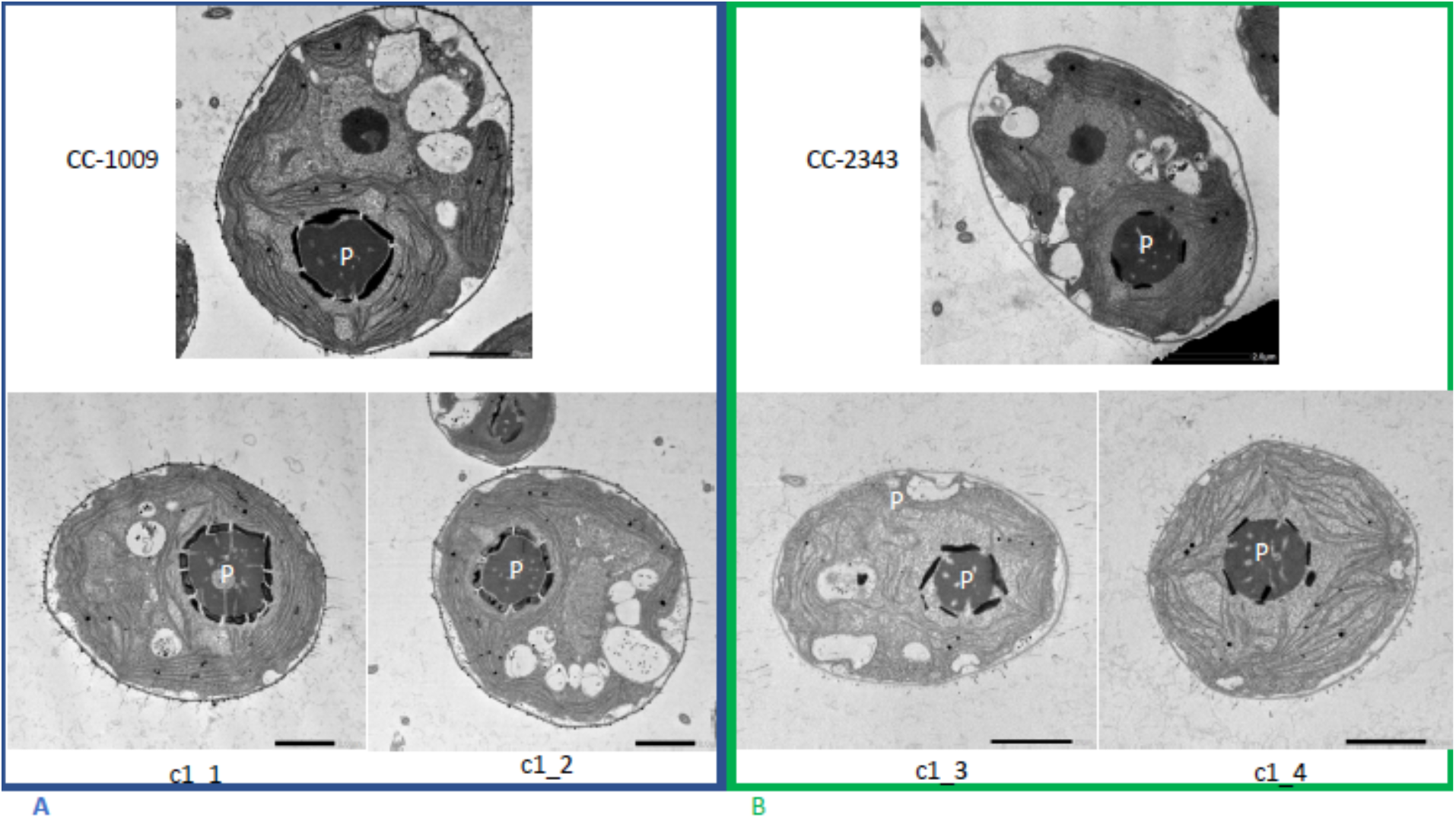
Representative TEM microscopy images of *Chlamydomonas* strains. Strains CC-1009 and CC-2343 are the parents and c1_1, c1_2, c1_3, c1_4 are the progeny. Images were taken 6 hours after the culture had been switched to from 5% to low (i.e. ambient) levels of CO_2_. Prior to switching the gas to ambient levels of CO_2_, cells had been grown in steady state conditions, with 5% CO_2_ with 14:10 hour (light:dark) sinusoidal illumination with peak light intensity of 2000 μmoles m^−2^ s^−1^, in minimal 2NBH media. Two of the progeny had sealed, continuous pyrenoids like CC-1009 (Panel A), while two others had fragmented pyrenoids resembling CC-2343 (Panel B). Cells here were fixed at 11:00 am, at 1945 μmoles m^−2^ s^−1^. Pyrenoids are labeled with “P.” Scale bar = 2 μm.

**Figure 8:**
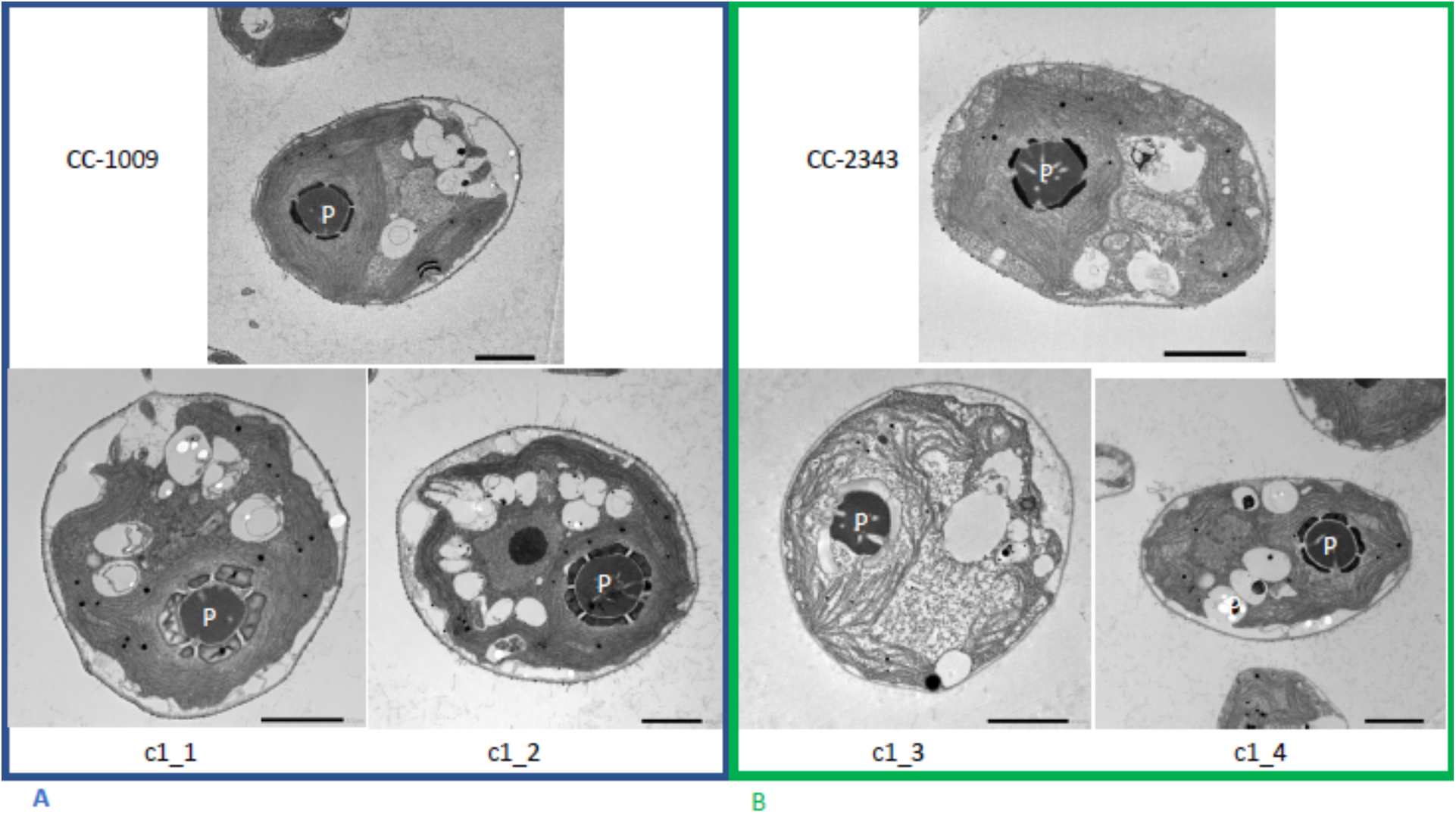
Representative TEM microscopy images of *Chlamydomonas* strains, the parents (CC-1009 & CC-2343), as well as their progeny c1_1, c1_2, c1_3, c1_4. Images taken 31 hours after the culture had been switched to from 5% to low (i.e. ambient) levels of CO_2_. Prior to switching the gas to ambient levels of CO_2_, cells had been grown in steady state conditions, with 5% CO_2_ with 14:10 hour (light:dark) sinusoidal illumination with peak light intensity of 2000 μmoles m^−2^ s^−1^, in minimal 2NBH media. At this time point, the differences in the pyrenoids were not as apparent between the two groups. Cells here were fixed at 11:00 am, at 1945 μmoles m^−2^ s^−1^. Pyrenoids are labeled with “P.” Scale bar = 2 μm.

### Pyrenoid formation is induced by exogenous and endogenously-produced H_2_O_2_, and inhibited by the ROS scavenger, ascorbic acid

The above results suggest that a product of photosynthesis common to high light, low CO_2_ and high O_2_ may trigger pyrenoid formation. As discussed below, one possible signal is H_2_O_2_. Figure 9 shows the effects of exogenous addition of H_2_O_2_ on the pyrenoid ultrastructure of *Chlamydomonas* parent lines. Cultures were harvested from photobioreactors in the morning (two hours after the start of illumination) and diluted by half with fresh minimal 2NBH media with 5mM bicarbonate – without (control) or with addition of 100 μM of H_2_O_2_. After six hours in low light (~85 μmol photons m^−2^ s^−1^), cells were fixed for EM as described in Materials and Methods. Strikingly, treatment with H_2_O_2_ resulted in the appearance of thick, well-sealed starch sheaths, for both CC-1009 (Figure 9A & 9B) and CC-2343 (Figure 9C & 9D). Morphometric analysis of the cells confirmed that there was a clear change in the morphology of the cells (Supplementary Material Figure 11). The hydrogen peroxide led to significant increases in the size of the pyrenoid, the prevalence of the starch sheath, and the appearance of the pyrenoid periphery mesh – which appears to cement the starch plates together.

**Figure 9:**
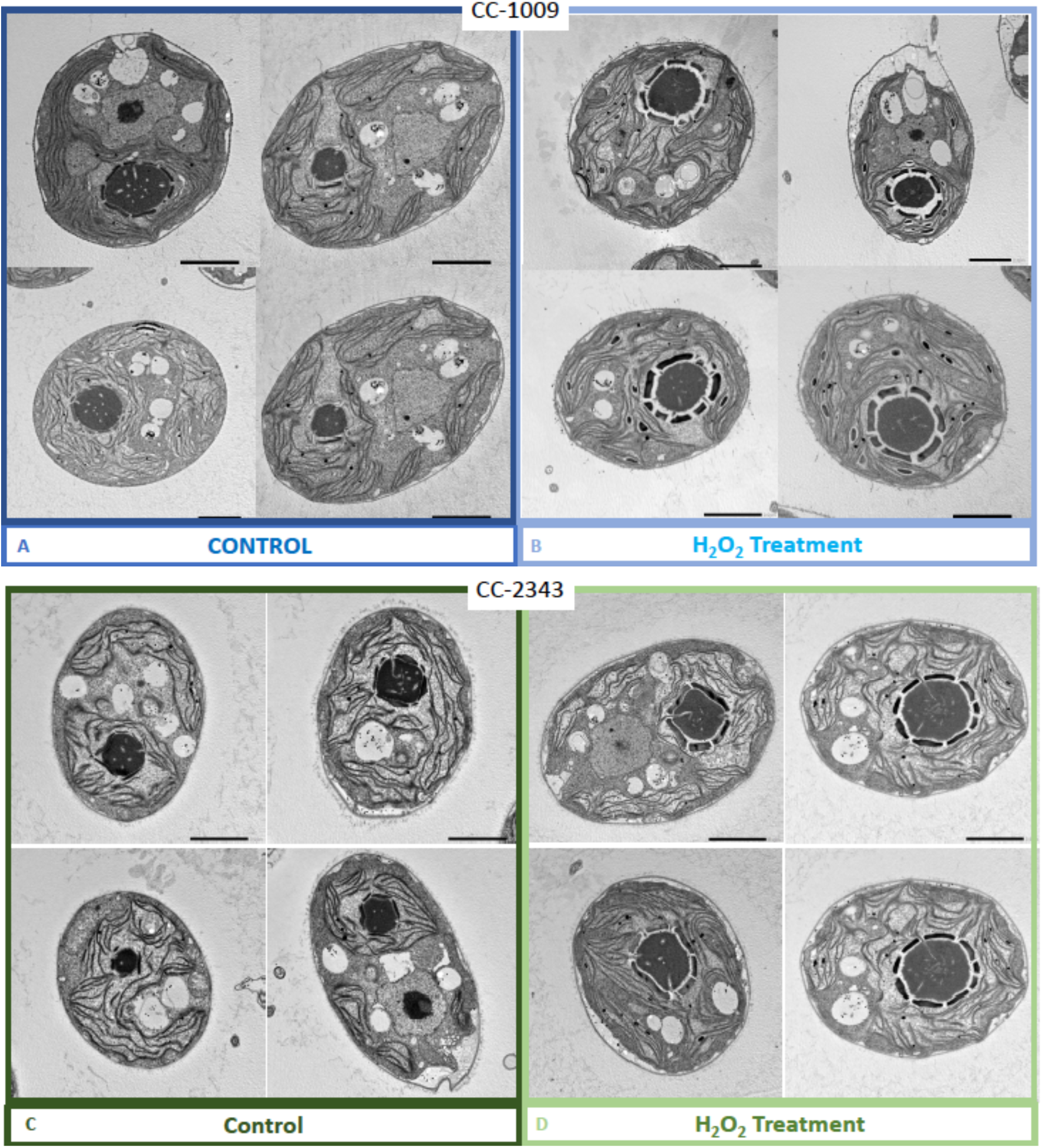
Representative TEM images of CC-1009 (Panels A & B) and CC-2343 (Panels C & D) control and cells treated, at 7:00 am in the morning, two hours after our sinusoidal light had turned on, with 100 μM of H_2_O_2_, and then exposed to 6 hours of low light (~85 μmoles photons m^−2^ s^−1^) with saturating (5mM) bicarbonate in minimal 2NBH media. Scale bar = 2 μm.

We also found that pyrenoids could also be induced in the presence of high bicarbonate via treatment with low concentrations of methyl viologen (Supplementary Material Figure 12) or metronidazole (Supplementary Material Figure 13), compounds known to induce internal hydrogen peroxide production by accepting electrons from PSI and passing them to O_2_, forming superoxide, which is converted to H_2_O_2_ by superoxide dismutase (Aksmann et al., 2016; Chang et al., 2013; Schmidt et al., 1977). The concentrations of these compounds did not inhibit growth or motility over the time scale of the experiment (~6 hours) and thus their effects are likely to be caused by ROS production or altered metabolic status rather than severe cell damage. Complementing these findings, treatment with two known H_2_O_2_ scavengers, ascorbic acid (Kuo et al., 2020; Nagy et al., 2015) (Supplementary Material Figure 14) or dimethylthiourea (Chang et al., 2013) (Supplementary Material Figure 15) prevented the formation of the pyrenoid under low CO_2_ conditions. Overall, these results are consistent with a role of H_2_O_2_ in triggering the formation of the pyrenoid, though it remains to be determined whether such effects are direct or indirect, e.g. resulting of altered metabolic status.

Hydrogen peroxide treatment was found also to affect the localization of rubisco (Figure 10), which is sequestered in the pyrenoid at low CO_2_ (Borkhsenious et al., 1998). We assessed changes in localization using a modified *Chlamydomonas reinhardtii* strain, CC-5357, expressing rubisco small subunit (Rbcs1) tagged with the Venus fluorescent protein (Mackinder et al., 2016). Under control conditions (5 mM bicarbonate, no H_2_O_2_ treatment), labelled rubisco was present throughout the chloroplast, with some localization in a pyrenoid matrix-like structure (Figure 10, Panel A). However, six hours after treatment with 100 μM H_2_O_2_, rubisco became strongly localized to the pyrenoid matrix (Figure 10, Panel B; Supplementary Material Figure 16), with very little fluorescent signal outside this structure (see quantification of fluorescence signal, Figure 10, Panel C). By contrast, no significant changes in rubisco localization were observed when upon addition of 100 μM H_2_O_2_ to TAP-grown cells (Supplementary Material Figure 17), implying that the effect was dependent on the photosynthetic state of the cells and/or suppressed in the presence of this organic carbon source. Consistent with this interpretation, cells grown on TAP plates showed no observable pyrenoid starch sheath by light microscopy or starch staining (Supplementary Material Figure 18) in contrast with what we observed with cells grown in liquid minimal media. Furthermore, when CC-5357 was grown on TAP plates, rubisco became completely dispersed throughout the stroma, with no evidence of a pyrenoid matrix-like structure (Figure 11; Supplementary Material Figure 19).

**Figure 10:**
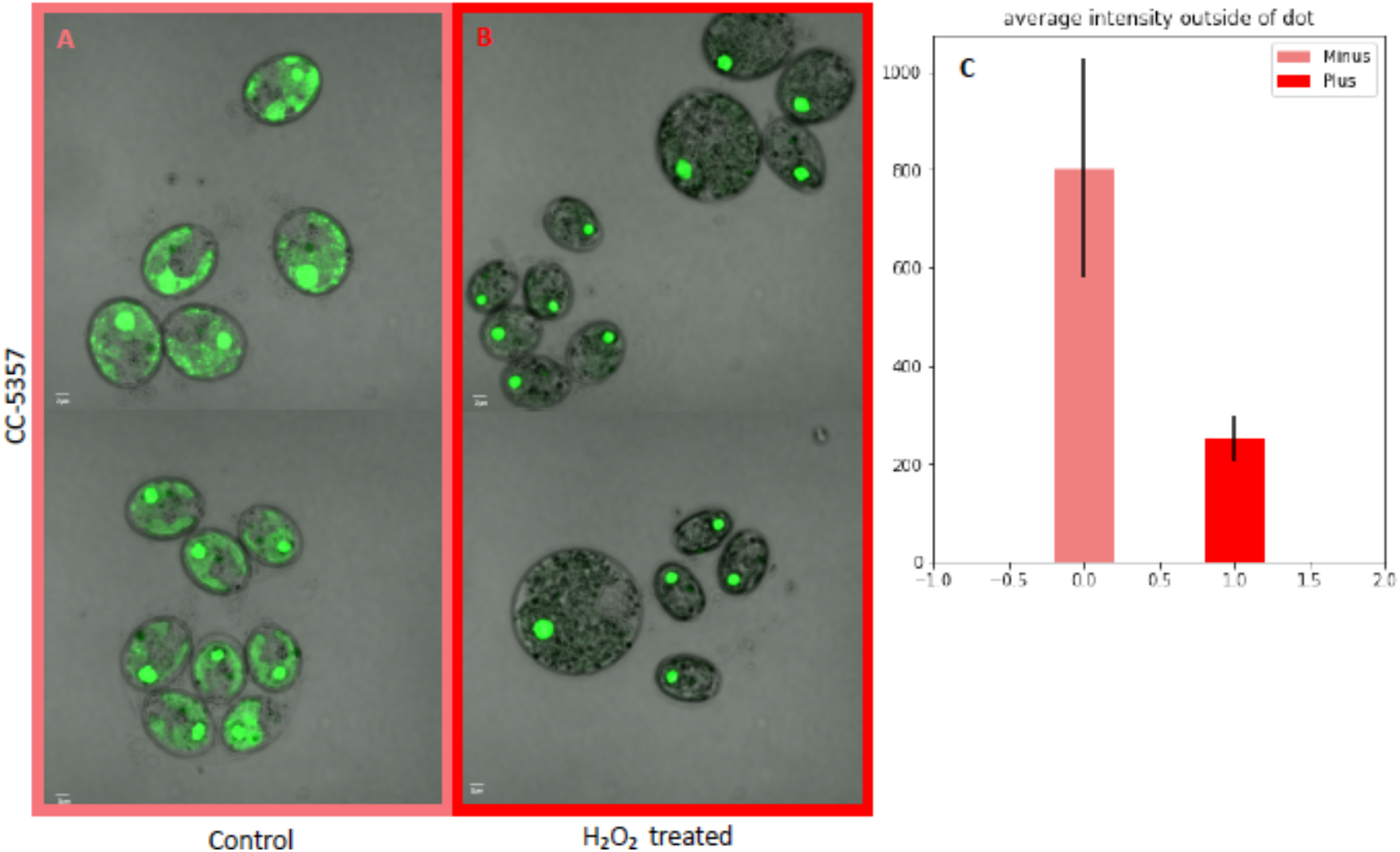
(Photos) Localization of rubisco determined by confocal microscopy of strain CC-5357, containing at RBCS1-Venus. (Bar Graph, Panel C) Average Intensity of fluorescent signal within a cell, outside of the pyrenoid region, without (**A**, minus) and with (**B**, plus) the addition of hydrogen peroxide. The average fluorescence intensity of the delocalized Venus Fluorescent Protein-labeled rubisco within *Chlamydomonas* cells was measured using the Olympus FluoView 1000 Advanced Software. For each cell measurement, a region encircling the *Chlamydomonas* cell membrane but excluding the pyrenoid was delineated and the average fluorescence intensity within the designated region was calculated. For each treatment, measurements were performed on approximately 20 cells from three separate areas, though more areas of cells were viewed to verify the consistency of the phenotype. Error bars on graph represent the standard deviation. Scale bar = 2 μm.

**Figure 11:**
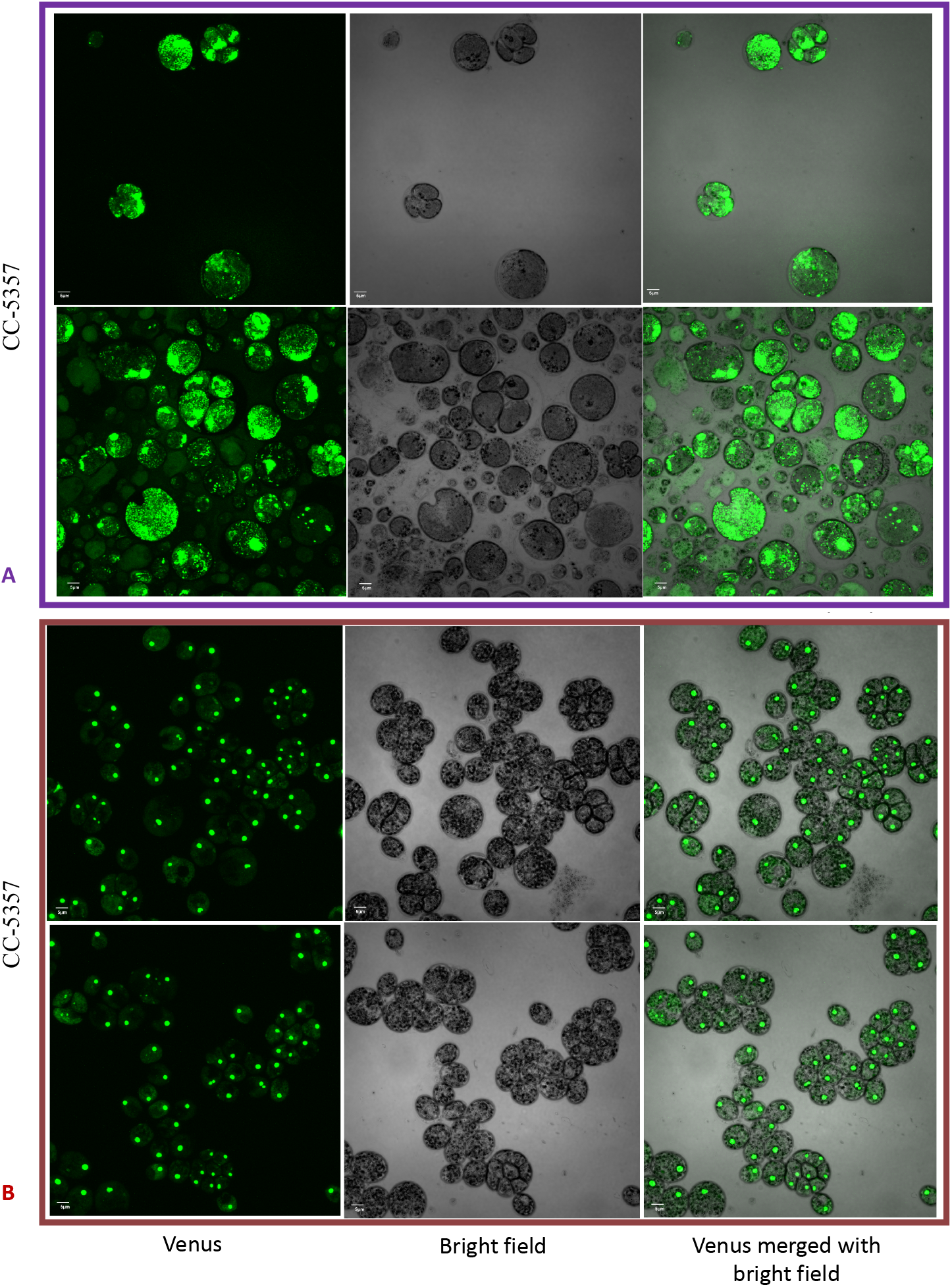
Confocal microscopy of CC-5357, which has a Venus labeled RBCS1, after being grown in on a TAP plate (Top, Panel A) showing rubisco completely de-localized and liquid TAP (bottom, Panel B), showing that rubisco has de-localized to some extent, but remains largely localized. Scale bar = 5 μm.

### H_2_O_2_ species is produced during exposure to hyperoxia

We next tested for differences in H_2_O_2_ production under hyperoxia in CC-2343 and CC-1009, using the Amplex™ Red Hydrogen Peroxide/Peroxidase Assay Kit (Invitrogen, Carlsbad, CA). We found that 6 and 31 h of exposure to hyperoxia resulted in a ~3-fold increase in H_2_O_2_ in CC-1009, but no significant changes in CC-2343 (Figure 12), though CC-2343 showed a somewhat higher basal level of H_2_O_2_ on a per cell basis. Figure 13 shows confocal laser-scanning microscope images of cells taken at steady state (Figure 13, Panels A & B) and at 31 hours hyperoxia (Figure 13, Panels C & D) and stained with 2’,7’-dichlorodihydrofluorescein diacetate (H_2_DCFDA), a general stain for reactive oxygen, sensitive to H_2_O_2_, singlet oxygen, superoxide, hydroxyl radical and various peroxide and hydroperoxides. Both cell lines accumulated ROS in response to hyperoxia. However, cells of CC-1009 showed accumulation of ROS that was highly localized in small structures (Figure 13, Panel D) consistent with peroxisomal microbodies (Lauersen et al., 2016). By contrast, CC-2343 cells showed weaker, more diffuse, staining throughout the cell, seeming to accumulate ROS throughout the thylakoids, which may be a result of rubisco inhibition, chloro-respiration, or superoxide formation (Figure 13, Panel C). We also observed, in CC-2343, cells uniformly stained with the H_2_DCFDA (Figure 13, Panel C), reflecting severe ROS accumulation/stress in CC-2343 in subpopulations of cells, stress that did not appear to occur in CC-1009.

**Figure 12:**
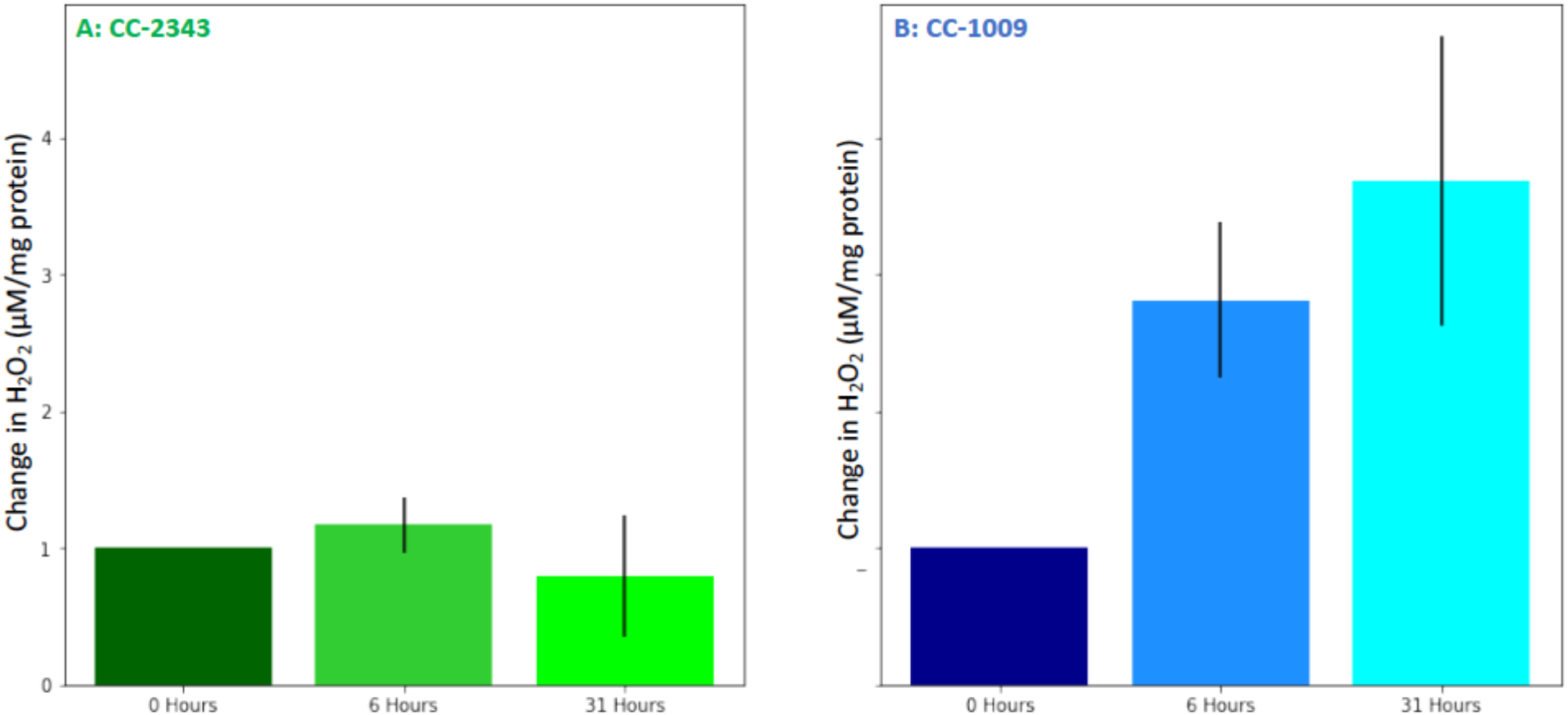
Changes in H_2_O_2_ in cellular extracts upon exposure to hyperoxia. Cells of CC-2343 (Panel A) and CC-1009 (Panel B) were rapidly broken and extracts assayed using the Amplex Red method just prior to (0 hrs) and at 6 and 31 hours exposure to 95% O_2_, 5% CO_2_, as described in Materials and Methods. Values shown are normalized to those taken at 0 hrs, when the values normalized to the extract’s protein contents were 3.37 μM for c2343 (Panel A) and .456 μM for CC-1009 (Panel B). Error bars represent the standard deviation among three biological replicates.

**Figure 13:**
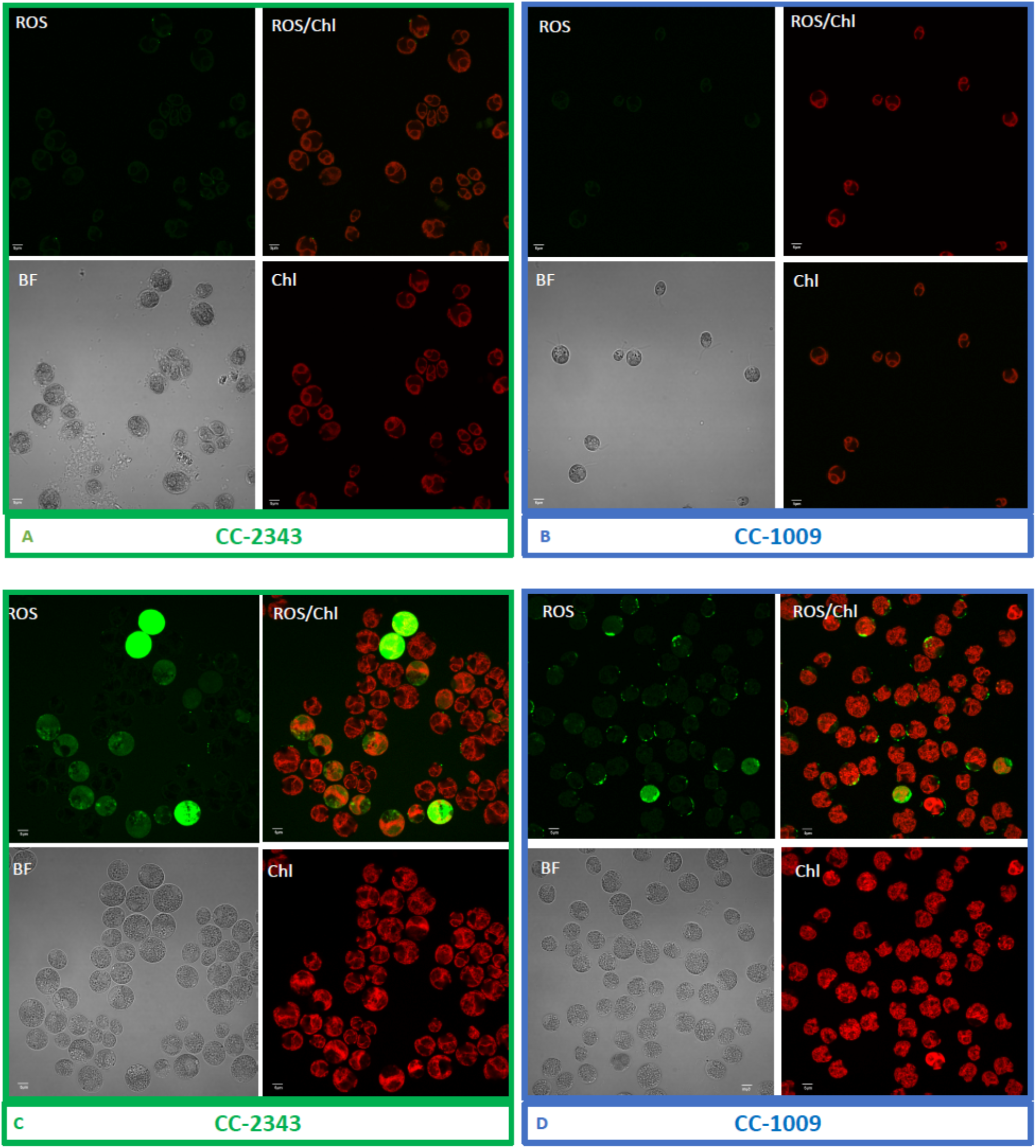
Confocal microscopy images of CC-2343 and CC-1009 showing ROS in cells growing at steady state (Panels A & B) and following exposure to 31 hours of hyperoxia (Panels C & D) of CC-2343 and CC-1009. H_2_DCFDA, a nonfluorescent probe that is converted into fluorescent dichlorofluorescein (DCF) by ROS, was used to detect the ROS. The ROS is indicated by the green fluorescence, while the auto- fluorescence of the chloroplasts is displayed in red. ROS, reactive oxygen species; Chl, chlorophyll; BF, bright field. Scale bars = 5 μmol.

### Cells pre-treated with exogenous H_2_O_2_ display higher oxygen compensation points

Figure 14 shows the effects of H_2_O_2_ pre-treatment on O_2_ levels in cell suspensions of CC-1009 and CC-2343 under saturating actinic illumination. In these experiments, we tested whether H_2_O_2_-induced formation of pyrenoids with tight sheaths allowed photosynthesis to occur at higher levels of O_2_. Prior to the traces, suspensions with 5 mM NaHCO_3_ were sparged with air to establish low dissolved O_2_ levels. At time zero, sparging was stopped and changes in dissolved O_2_ were monitored with a luminescence-based O_2_ sensor (see Materials and Methods). The initial rise in O_2_ reflects when the rate of net assimilation was maximal, under conditions when inorganic C supply was replete (5 mM HCO_3-_) and O_2_ levels were low. These slopes were within 15% of one another for both control and H_2_O_2_-treated CC-1009 (127.7 and 142.7 μM O_2_ min^−1^) and CC-2343 (104.28 and 109.64 μM O_2_ min^−1^) suspensions. After about 20 minutes, the rise in O_2_ levels slowed, eventually reaching quasi-steady-state levels that represented the “oxygen compensation point” where O_2_ evolution from PSII was counterbalanced by O_2_ uptake. Switching off the actinic light at ~57 minutes led to O_2_ uptake, the initial rate of which likely represents the gross O_2_ uptake, which is counterbalanced by O_2_ evolution. Nearly equal during steady-state illumination, the two canceled each other out during the periods of light exposure. For control cells, the O_2_ compensation points (the O_2_ levels when the rate of O_2_ uptake balanced that of evolution) for CC-2343 and CC-1009 were approximately 1070 and 1227 μM O_2_ (P<0.05), respectively, implying, because it reaches the compensation point at a higher O_2_ level, that CC-1009 was able to more effectively select for CO_2_ uptake over O_2_ reduction. The rates of uptake of O_2_ after illumination were slightly slower in CC-2343 (−36.4 μM O_2_ min^−1^) than CC-1009 (−44.3 μM O_2_ min^−1^) indicating that the lower compensation point was caused by a combination of decreased linear electron flow and increased O_2_ uptake. Strikingly, pre-treatment with H_2_O_2_ led to significant (P<0.05) increases in the O_2_ compensation points for both CC-2343 and CC-1009, to about 1233 and 1356 μM O_2_ min^−1^ respectively, confirming that treated cells were better able to discriminate between uptakes of CO_2_ and O_2_. After the cells reached (essentially) steady-state levels of O_2_, the actinic light was switched off. The initial slopes of O_2_ uptake were then measured (by fitting the decay to pseudo-first order decay kinetics and taking the initial rate), to give an estimate of the rates of O_2_ evolution and uptake. CC-1009 cells showed similar O_2_ uptake slopes in treated H_2_O_2_-treated (45.9 μM O_2_ min^−1^) and untreated (44.3 μM O_2_ min^−1^) suspensions, likely indicating that the rates of electron flow were also similar, but that the preferential fluxes of electron into assimilation allowed for a greater accumulation of O_2_ in the treated cells. By contrast, CC-2343 cells showed a significant increase in the initial slopes of O_2_ consumption in the the treated (41.99 μM O_2_ min^−1^) compared to control (36.4 μM O_2_ min^−1^) suspensions, suggesting that the high O_2_ levels suppressed overall rates of linear electron flow (LEF) in the untreated cells (Supplementary Material Table 3).

**Figure 14:**
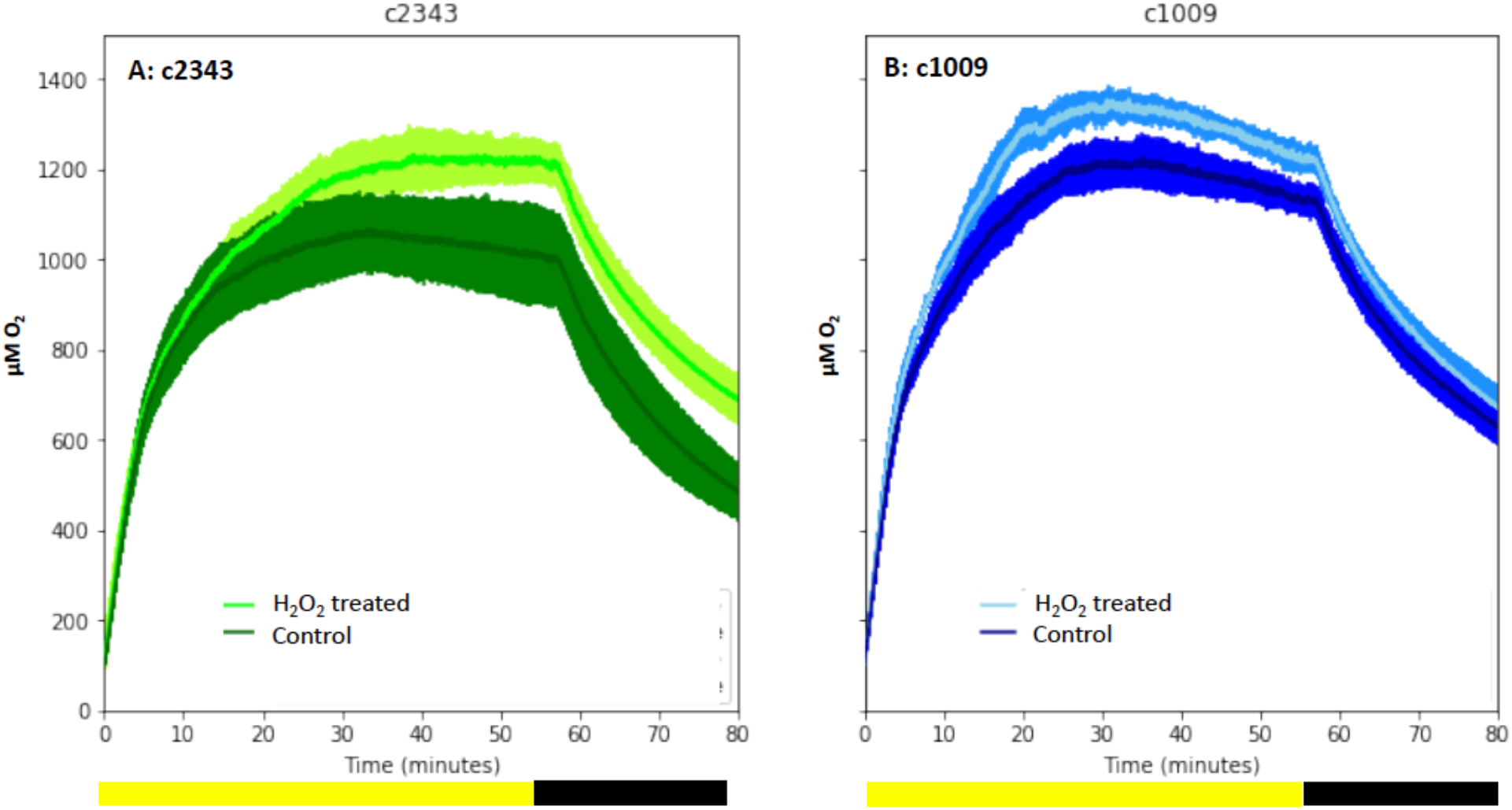
Oxygen evolution of strain CC-2343 (Panel A) and CC-1009 (Panel B) with and without the pre-treatment of 100 μM H_2_O_2_. Shading represents 95% confidence intervals between the three biological replicates for each treatment. At approximately 3500 seconds (denoted by yellow bar on x axis), the light was turned off denoted by black bar on x axis).

## DISCUSSION

### The induction of pyrenoid biosynthesis under hyperoxia and the role of H_2_O_2_

Photosynthesis can be inhibited by one of its major products, molecular oxygen. This is known to occur in certain aqueous environments, such as when algae ponds are enriched with CO_2_ and photosynthesis can proceed rapidly, but diffusion of O_2_ is slow, leading to super-saturated oxygen levels, which can feedback limit productivity (Livansky, 1996; Pulz, 2001; Raso et al., 2012; Torzillo et al., 1998; Vonshak et al., 1996; Weissman et al., 1988). Little is known about the physiological impact of hyperoxia or the mechanisms by which some species algae are able to ameliorate its effects. Here, we took advantage of an observation that two strains of *C. reinhardtii* – and their meiotic progeny – showed distinct tolerances to hyperoxia to probe such adaptations.

Most previous work on the pyrenoid has focused on its activation by low CO_2_ and the its role in the CCM (Borkhsenious et al., 1998; Freeman Rosenzweig et al., 2017; Mackinder et al., 2017; Ramazanov et al., 1994). Both parent lines in our study and all progeny showed low CO_2_-dependent pyrenoid formation. When cells were sparged with 5% CO_2_ for one minute every hour and under normoxia (~21%), the pyrenoid starch sheaths dissociated at night. This result was consistent with previous observations which demonstrated that starch formation around the pyrenoid is correlated with light and the state of the CCM (Borkhsenious et al., 1998; Kuchitsu et al., 1988; Lin and Carpenter, 1997; Ramazanov et al., 1994). Also consistent with the cited previous work, low-light with mixotrophic conditions resulted in rubisco delocalization. However, this is the first study, to our knowledge, to show that the most complete degree of rubisco delocalization occurs when an algae is grown, rather than aquatically in liquid TAP, on a TAP agar plate exposed to air (Figure 11; Supplementary Material Figure 19). At high light, pyrenoids were formed even when CO_2_ or bicarbonate levels were maintained at high levels (Figure 6), agreeing with previous assertions that light plays a role in pyrenoid biosynthesis (Kuchitsu et al., 1988; Lin and Carpenter, 1997).

Strikingly, we also observed strong pyrenoid formation, with especially tight, thick and well-sealed starch sheaths in CC-1009, c1_1, and c1_2 under hyperoxia (95% O_2_), despite the high CO_2_ levels (Figures 1 & 2). One possible explanation for this observation is that an in-common by-product of photosynthesis and photorespiration under low CO_2_ and hyperoxia acts to induce pyrenoid formation. Hydrogen peroxide is an obvious candidate for such a role because it is well-documented to act as a signal molecule (Foyer et al., 2009) and its production is increased under high light (Roach et al., 2015), high O_2_ (Kim and Portis, 2004) or low CO_2_ (Foyer et al., 2009). H_2_O_2_ is a product of the light reactions, through an alternative electron acceptor pathway such as the Mehler peroxidase reaction (MPR) or the water-water cycle, which is expected to be more active under conditions when light input exceeds the capacity of assimilation (Badger et al., 2000; Mehler, 1951; Strizh, 2008). H_2_O_2_ can also be produced as a by-product of photorespiration (Kim and Portis, 2004). *Chlamydomonas* possesses two pathways for oxidation of glycolate during photorespiration, one involving glycolate oxidase in the peroxisome, which uses O_2_ as an electron acceptor and produces H_2_O_2_, and another involving glycolate dehydrogenase (GLYDH) in the mitochondrion, which uses ubiquinone as an electron acceptor and presumably does not produce H_2_O_2_ (Janssen et al., 2014). A reasonable explanation is that, under conditions of high light, low CO_2_ or high O_2_ production of H_2_O_2_ by the glycolate pathway can act as a signal to induce pyrenoid formation.

We found that, in autotrophically-grown cells, exogenous addition of H_2_O_2_ in the presence of light strongly induced within 6 hours the formation pyrenoid starch sheaths (Figure 9; Supplementary Material Figures 20 & 21), and caused rubisco to localize into the pyrenoid (Figure 10). Regardless of which strain was used, the starch sheaths formed after addition of H_2_O_2_ had tight, thick structures in both parent lines, though CC-1009 seemed to still display slightly more robust starch plates (Figure 9). These structural changes were accompanied by increased O_2_ compensation points (Figure 14), indicating an increased ability to discriminate between O_2_ and CO_2_ as O_2_ levels increased. Our working hypothesis is that H_2_O_2_-induced formation of pyrenoids with tight sheaths allowed the accumulation of higher concentrations of CO_2_ at the active site of rubisco, outcompeting or shielding out higher levels of O_2_. Further, the formation of these pyrenoids enhance the discrimination of CO_2_/O_2_, implying that H_2_O_2_ induction of pyrenoids could convey performance advantages under hyperoxia. Consistent with this hypothesis, the induction of the CCM has been found to be coordinated with that of genes for photorespiratory enzymes, though the specific metabolic control of this co-regulation had remained unknown (Tirumani et al., 2019). Interestingly, a separate RNA expression study (Blaby et al., 2015) did not show strong induction of pyrenoid components by H_2_O_2_, but, importantly, was conducted on TAP-grown cells, which we found do not show H_2_O_2_-induced formation of the pyrenoid (Supplementary Material Figure 17).

Debate remains about the signal that induces the CCM (Spalding, 2009; Spalding et al., 2002; Vance and Spalding, 2005), which consists not only of the pyrenoid but also the inorganic carbon transporters (Spalding, 2008) and carbonic anhydrases (Moroney et al., 2011); some have argued that the signal is CO_2_ itself, while others have proposed that the signal is a metabolite produced under low CO_2_ during photosynthesis or photorespiration. The later, termed the ‘metabolic signal hypothesis,’ (Spalding, 2009) proposed that photorespiratory intermediates could serve as a trigger for CCM induction (Marcus et al., 1983; Suzuki et al., 1990). The hypothesis was rooted in observations that, unlike wild type cells, various photosynthetic mutants did not exhibit CCM activity under low CO_2_, and that CCM induction in wild type cells required light (Dionisio et al., 1989a, b; Dionisio-Sese et al., 1990; Spalding and Ogren, 1982; Spencer et al., 1983; Tirumani et al., 2014; Villarejo et al., 1996). Also implying that other factors, apart from CO_2_, played a role in CCM induction, decreased O_2_ tension and photorespiratory inhibitors, in low CO_2_ conditions, also decreased carbonic anhydrase induction (Ramazanov and Cardenas, 1992; Spalding and Ogren, 1982; Villarejo et al., 1996),

Bozzo et al (Bozzo et al., 2000) argued against the metabolite hypothesis in *Chlorella*, based on observations that: 1) photorespiratory inhibitors, which should result in an accumulation of photorespiration intermediates, failed to induce the CCM under high CO_2_ and 2) the expression of transcripts for a subset of carbonic anhydrases increased under low CO_2_ even in the dark (though to a lesser extent than in the light). Similar results have been found in several *Chlorella* species (Matsuda and Colman, 1995; Shiraiwa and Miyachi, 1985; Umino et al., 1991), suggesting that the induction of at least some CCM components can occur in the dark. However, it is unclear how relevant these results are, considering that the pyrenoid is not formed in the dark (Kuchitsu et al., 1988; Lin and Carpenter, 1997). It was also found in *Chlamydomonas* that changing O_2_ levels (from 2-20%) did not affect growth, photosynthetic rate, or the induction of periplasmic carbonic anhydrase (Cah1) or glycolate dehydrogenase (Gdh) genes, over a wide range of CO_2_ levels (Vance and Spalding, 2005). It is worth noting, though, that none of these previous experiments were conducted under true hyperoxia (O_2_ levels above partial pressure of 21%), where we observe strong induction of the pyrenoid, and thus they do not exclude product signaling under our observed conditions.

There remains the possibility of multiple signals for the CCM. There is differential regulation of low CO_2_ induced polypeptides in *Chlamydomonas*, with some only being induced in the light, while for others light is not necessary (Villarejo et al., 1996). Also, the observation that there are multiple acclimation states, with some mutants tolerant to very low CO_2_ but not low CO_2_, suggest the existence of multiple types of signaling (Spalding et al., 2002). Our findings that a key aspect of the CCM, the pyrenoid, can be induced, even under high CO_2_ (i.e. with hyperoxia and H_2_O_2_) disproves, to our knowledge for the first time, that low CO_2_ is a necessary condition for any aspect of CCM induction. Our results lead us to propose that H_2_O_2_, a by-product of photosynthesis, particularly under low CO_2_ and high O_2_, may fulfill the previous proposed ‘metabolic’ signal. Hydrogen peroxide is widely known to be a signal for a variety of stress related responses (Blaby et al., 2015; Zalutskaya et al., 2019), and has been found to alter the state of redox homeostasis in *Chlamydomonas* (Pokora et al., 2018). It has also been assigned roles in regulating a range of photosynthetic and associated processes in plants and algae (Berens et al., 2019; Foyer and Noctor, 2009), particularly those related to responses to CO_2_ levels and the induction of photorespiration (Foyer et al., 2009). Interestingly, in higher plants, H_2_O_2_ has been suggested to play a role in the response to varying levels of CO_2_ (Foyer et al., 2009). For example, *Sorghum* (C4) plants grown under conditions with lower amounts of photorespiration (i.e. elevated CO_2_) have decreased thickening of the bundle sheath cells (Watling et al., 2000), which, since they restrict the diffusion of CO_2_ out of bundle sheath cells and thereby allow for efficient capture by rubisco, can be interpreted as structures analogous to the starch sheath of the pyrenoid.

Hydrogen peroxide has also been implicated in regulating cyclic electron flow (CEF) in vascular plants, both by inducing the expression of the thylakoid Complex I (or NDH) (Casano et al., 2001; Gambarova, 2008) and by activation of existing enzymes (Strand et al., 2015). It is not known, however, if H_2_O_2_ acts directly as a signaling agent, or indirectly, e.g. by altering the activities of assimilatory enzymes (Strand et al., 2015) possibly through redox balancing enzymes such as the peroxiredoxins (Vaseghi et al., 2018). CEF is thought also to play central role in providing the energy needed to power CCMs, including that in *Chlamydomonas* (Lucker and Kramer, 2013). Our findings indicating that H_2_O_2_ may be the signal for the synthesis of a central component of the CCM, the pyrenoid, suggests that a common molecular by-product of photorespiration can set off a coordinated response; inducing the formation of the pyrenoid and also the metabolic processes to power the pumping of bicarbonate into it.

It has been argued that mixotrophic conditions alter the relationship between the onset of photorespiration and the expression of the CCM (Tirumani et al., 2019), and that photorespiration, hydrogen peroxide detoxification, and acetate assimilation (i.e. the glyoxolate cycle) are all localized in the peroxisomal microbodies (Lauersen et al., 2016). In this regard, it is intriguing that ROS labeling under hyperoxia was strongly localized in CC-1009 but not CC-2342 (Figure 13), hinting that H_2_O_2_ produced in a specific subcellular location and process may play a role in the differential development of the pyrenoid in the two parent lines, as discussed below. Taken together, these data sets are consistent with control of pyrenoid morphology at multiple levels, perhaps similar to the processes that regulate the expression of LHCSR3, involved in photoprotective nonphotochemical quenching, which is regulated by light quality and CO_2_ availability (Maruyama et al., 2014; Semchonok et al., 2017).

### Possible linkages between H_2_O_2_ signaling, pyrenoid morphology and natural variations in responses to hyperoxia

By comparing genetically distinct isolates and their progeny, one can explore possible mechanistic bases for responses to hyperoxia. We present here data from a limited set of progeny, which nevertheless reveals a segregation pattern which allows us to test certain future hypotheses. A more detailed analysis of a large number of progeny will be presented elsewhere. The most striking differences we observed between the lines were in the morphology of the pyrenoids (Figure 1, Figure 2), with the hyperoxia tolerant lines (CC-1009, c1_1, c1_2) showing thick, tightly sealed starch sheaths, while the sensitive lines (CC-2343, c1_3, c1_4) tending to have pores or gaps in the starch sheaths (Figures 1-3), suggesting that structural/functional differences in these sub-organelle compartments may play a role in the distinct responses to high O_2_. These differences appear to be most obvious during hyperoxia, and all lines showed disappearance of the pyrenoid structures under high CO_2_/low light (Figures 5). Most interestingly, exogenous H_2_O_2_ led to synthesis of thick, tight pyrenoids in all lines, implying that the distinct morphologies is caused at least in part from differences in signaling, rather than structural components.

Given the possibility that H_2_O_2_ acts as a signal for pyrenoid biosynthesis, we tested for differences in its production under hyperoxia. While both parent lines showed increased whole-cell H_2_O_2_ production as measured by the Amplex assay (Figure 12), the localization of ROS production assessed by H_2_DCFDA fluorescence in confocal microscopy showed distinct localization patterns, with CC-1009 showing strongly localized dye fluorescence (Figure 13, Panels B & D) whereas CC-2343 showed diffuse staining throughout the cell (Figure 13, Panels A & C). Because the H_2_DCFDA is a general ROS indicator, it is not possible to unambiguously identify the specific reactive oxygen species, but one possible interpretation is that different localization patterns reflect the mechanism of ROS formation. The localized staining in CC-1009 is consistent with H_2_O_2_ produced in the peroxisome through photorespiration. By contrast, in CC-2343, the diffuse staining may reflect a range of different ROS, including but not limited to H_2_O_2_, ^1^O_2_ and O_2_·^-^, produced by excitation of the light reactions and other processes (Badger et al., 2000).

We have several hypotheses regarding why the two lines may have differences in the signaling and formation of their pyrenoid starch sheaths. One is that there might be variations in the strains’ utilization of the alternative photorespiration route that uses the glycolate dehydrogenase (GLYDH) enzyme, a route which does not result in hydrogen peroxide formation (Janssen et al., 2014).

Similarly, in the future we will investigate how the pyrenoid ameliorates the stresses of hyperoxia, with possibilities beyond photorespiration. Our rubisco assays (Supplementary Material Figure 3) suggest that increased O_2_ fixation or ROS production may lead to greater inhibition of rubisco in CC-2343 compared to CC-1009, possibly leading or concomitant to a general breakdown of the cell’s machinery, as evidenced by the lower autotrophic grow rates of CC-2343 at high O_2_ (Figure 3) and lower rates of oxygen evolution (Figure 14). Such a model is also consistent with the diffuse ROS staining observed in CC-2343, as the mismatch in light input and downstream assimilatory capacity could result in the accumulation of not just H_2_O_2_, but also ^1^O_2_ and O_2_·^-^ (Peng et al., 2017), forms of ROS that may reflect high levels of oxidative damage.

### Eco-physiological Implications

For over a hundred years it has been known that *Chlamydomonas* strains show distinct pyrenoid structures (Pasher, 1918), though the physiological implications of these natural variations remain poorly understood. A few studies have noted structural differences in pyrenoid starch sheaths, and linked these differences to environmental CO_2_ or organic carbon availability (Morita et al., 1998, 1999; Nozaki et al., 1994).

As discussed above, it is well established that the pyrenoid can allow algae to overcome critical limitations of low CO_2_ levels often encountered in aqueous environments. However, under very high CO_2_ levels, which are also encountered in certain environments, the sequestering of rubisco into the pyrenoid may impose rate limitations, or additional energy requirements, at the level of pumping of bicarbonate. Also, when rubisco is outside of the pyrenoid, it is thought that more of its surface area is exposed and its catalytic rate increases (Badger et al., 1998). A fragmented starch sheath may more easily allow migration in and out of the pyrenoid matrix. In two species of *Gonium*, the species with the more porous starch sheaths exhibited a higher ratio of rubisco migrating out of the pyrenoid in response to the addition of sodium acetate (Nozaki et al., 1994). Among closely related *Chlamydomonas* and *Chloromonas* strains, those with tight (which were termed “typical”) pyrenoids were able to accumulate higher levels of inorganic carbon when CO_2_ was low compared to those with fragmented or porous (termed “atypical”) pyrenoid starch sheaths (Morita et al., 1999). On the other hand, *Chloromonas* species closely related to *Chlamydomonas* but lacking pyrenoids showed higher rates of max O_2_ evolution when grown under elevated CO_2_ (Morita et al., 1998), which could be attributed to the greater accessibility of rubisco to diffusible CO_2_.

Some algae lack pyrenoids altogether, and are found in environments expected to have high CO_2_ and low or atmospheric oxygen levels. For example, *Coccomyxa*, an aerial grown lichen photobiont, completely lacks pyrenoids (Palmqvist et al., 1994; Palmqvist et al., 1995). Compared to that in *Chlamydomonas*, *Coccomyxa* prefers CO_2_ as a substrate over HCO_3_, similar to C3 plant cells (Palmqvist et al., 1994; Palmqvist et al., 1995). It is important to note, though, that exposure to air allows for rapid diffusion of O_2_: Even high rates of photosynthesis in *Coccomyxa* will not result in hyperoxia. In light of these studies, it seems fitting that CC-2343 and the progeny with porous pyrenoids grew better on a TAP plate exposed to air, and that rubisco most freely distributes through the chloroplast in *Chlamydomonas* when grown mixotrophically exposed to air, rather than aquatically (Figure 11; Supplementary Material 19).

By contrast, green algae can generate strongly hyperoxic conditions specifically when inorganic carbon is plentiful. Our demonstration that pyrenoids are induced under these conditions suggests that they can function, in addition to overcoming slow assimilation when CO_2_ is limiting, in preventing damage caused by high levels of the product O_2_. Inducing the CCM should both increase the concentration of CO_2_ above its K_M_ at rubisco and outcompete O_2_ at the rubisco active site. Higher O_2_ levels (under hyperoxia) will require correspondingly higher local CO_2_ levels, in turn requiring tighter diffusional barriers to the escape of CO_2_ from the pyrenoid (Wang et al., 2015; Yamano et al., 2015). It is also possible that the tight starch sheaths will partially block O_2_ from diffusing into the pyrenoid, and if the uptake of O_2_ by rubisco is faster than its replacement by diffusion across the sheath, such a barrier could effectively decrease the O_2_ levels in the matrix.

### Conclusions

The work presented above leads us to propose that, under combinations of light, high O_2_ and/or low CO_2_, the production of H_2_O_2_ becomes elevated, activating the formation of the pyrenoid and thickening of the starch sheath, leading to the classical response that allows cells to better discriminate between CO_2_ and O_2_ (Aizawa and Miyachi, 1986; Badger et al., 1980; Borkhsenious et al., 1998; Manuel and Moroney, 1988; Ramazanov et al., 1994; Spalding et al., 1983). We demonstrate that the pyrenoid, a key component of the algal CCM, can be induced under high CO_2_, by hyperoxia or H_2_O_2_. Our results strengthen the ‘metabolite signaling hypothesis,’ (Spalding, 2009), which can explain the regulation of pyrenoid formation by multiple photosynthetic conditions, including CO_2_, O_2_, and its light dependence. Our results further suggest that differences in this signaling contribute, at least in part, to the observed natural varaition in pyrenoids (Pasher, 1918) as well as tolerances to hyperoxia. Several open questions remain, including whether a H_2_O_2_ signal works alone or in conjunction with a CO_2_ signal for some aspects of the CCM, the precise nature and scope of the H_2_O_2_ response, the biochemical and genetic components involved, and whether more robust pyrenoid structures, by themselves, can improve growth under hyperoxia.

