## Supplemental Information for "The Induction of Pyrenoid Synthesis by Hyperoxia and its Implications for the Natural Diversity of Photosynthetic Responses in *Chlamydomonas*"

**SUPPLEMENTARY MATERIAL**

**Supplementary Material Table 1**: 2NBH Growth Media

| Volume of Stock Solution (ml/l) | Stock Solution | Stock Solution Content (grams) |
|---|---|---|
| 2 | NaNO3 | 100 g/400 ml |
| 1 | $CaCl_2$ | 10 g/ 400 ml |
| 1 | $MgSO_4$ $7H_2O$ | 30 g / 400 ml |
| 1 | TAP Phosphate Solution | 28.8 g $K_2HPO_4$ + 14 $KH_2PO_4$ |
| 1 | NaCl | 10 g NaCl / 400 ml |
| 1 | Hutner Solution | |

**Supplementary Material Table 2**: Antibodies used in Western Blotts.

| Protein | Component | Comment |
|---|---|---|
| PsbA | PSII | D1 protein of PSII, C-terminal |
| PsbO | PSII | oxygen evolving complex of PSII |
| Lhcb2 | LHC | LHCII type II chlorophyll a/b binding protein |
| Lhcb6 | LHC | LHCII chlorophyll a/b binding protein CP24 |
| Lhca2 | LHC | PSI type II chloropyll a/b-binding protein |
| PsaC | PSI | PSI-C, subunit of PSI |
| PsaD | PSI | PSI-D, subunit of PSI |
| PsaH | PSI | PSI-H subunit of photosystem I, Chlamydomonas |
| Cyt f | Cyt b6f | cytochrome f subunit of cytochrome b6f complex |
| Rieske | Cyt b6f | Rieske iron-sulfur protein of cytochrome b6f complex |
| ATPase | ATP Synthase | ATP synthase, whole enzyme |
| Rubisco Lg | Stroma | RbcL \| Rubisco large subunit, form I and form II |

**Supplementary Material Table 3**: Rates of oxygen evolution in CC-2343 and CC-1009 (µM $O_2$ $min^{-1}$) and maximum oxygen compensation point in control and cells pre-treated for 6 hours with hydrogen peroxide (see Figure 14 for graphs).

| | Initial slope | 15 minutes slope | 58 minutes slope | 65 minutes slope | Max $O_2$ |
|---|---|---|---|---|---|
| **CC-2343 with $H_2O_2$** | 109.64 | 19.52 | -41.99 | -24.57 | 1233.27 |
| **CC-2343** | 104.28 | 11.12 | -36.39 | -23.67 | 1069.67 |
| **CC-1009 with $H_2O_2$** | 142.68 | 30.79 | -45.91 | -25.066 | 1356.18 |
| **CC-1009** | 127.68 | 23.88 | -44.31 | -20.99 | 1227.82 |

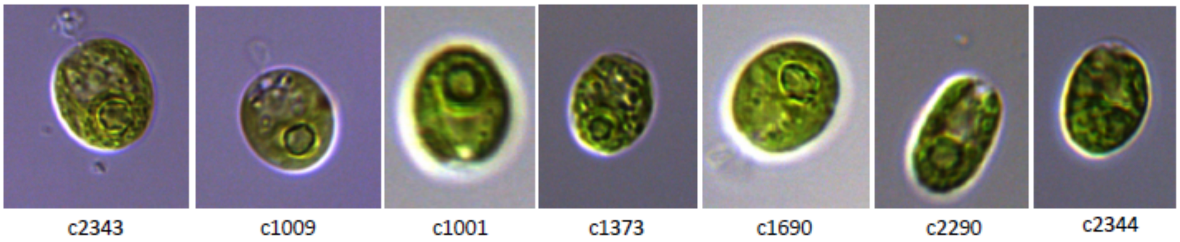


**Supplementary Material Figure 1**: Exemplary pictures of ecotypes which were examined in this study showing various pyrenoid formations, with CC-1009 and CC-1001 having the most clear "closed" type. Cultures were grown within flasks in minimal (2NBH) media under 50 µmoles $m^{-2}$ $s^{-1}$ of PAR.

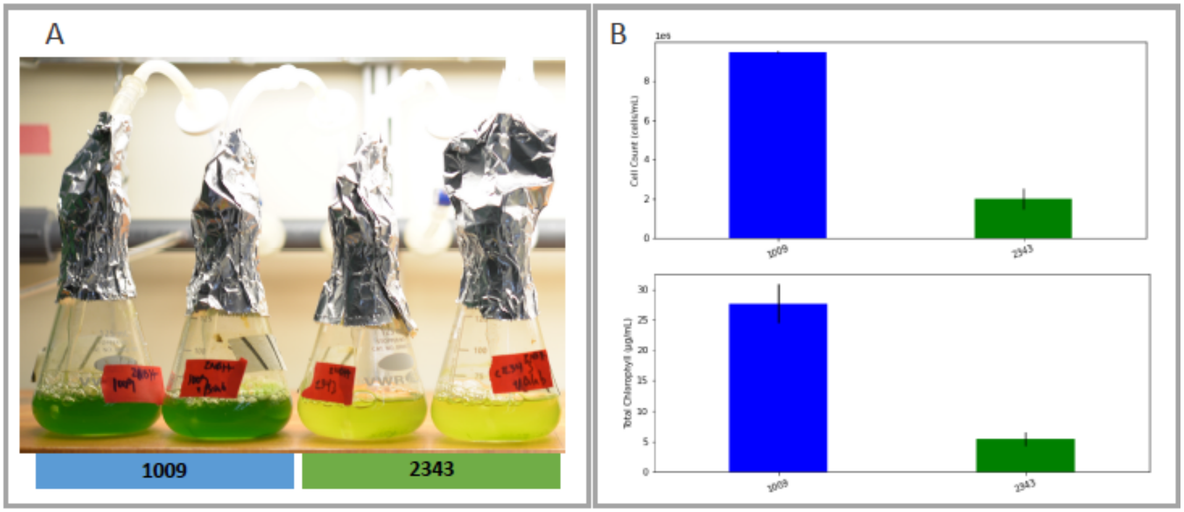


**Supplementary Material Figure 2**. Panel A) Photo of five days after 50ml of fresh 2NBH media was inoculated to $1x10^5$ cells/ml, and cultures were continuously bubbled with 5% $CO_2$ and 95% $O_2$. Panel B) Graph of cell counts and chlorophyll content. Error bars represent the standard deviation of the two biological replicates.

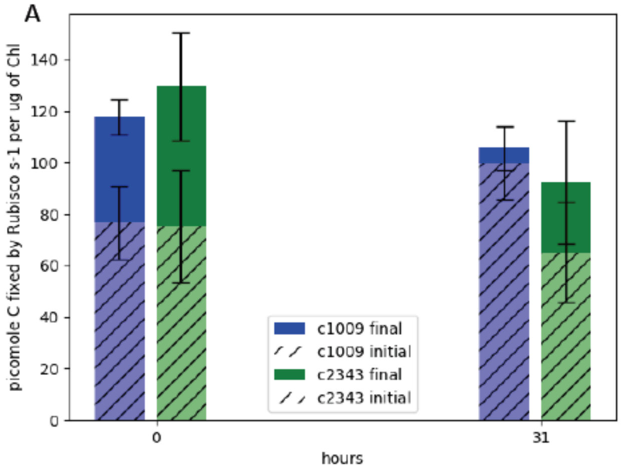


B

| | 0 Hours | | | | | 31 Hours | | | | |
|---|---|---|---|---|---|---|---|---|---|---|
| | Initial | STD (+/-) | Final | STD (+/-) | % Activity | Initial | STD (+/-) | Final | STD (+/-) | % Activity |
| CC-1009 | 76.70 | 14.2 | 117.73 | 6.86 | 65.15% | 99.75 | 14.31 | 105.58 | 8.39 | 94.48% |
| CC-2343 | 75.24 | 21.8 | 129.66 | 20.9 | 58.03% | 65.24 | 19.35 | 92.44 | 23.96 | 70.58% |
| P-Values | 0.915 | | 0.322 | | | 0.068 | | 0.421 | | |

**Supplementary Material Figure 3**: Panel A) Graph of effects of hyperoxia on activity of rubisco in CC-1009 and CC-2343. Raw extracts of the cells prior to (zero hours) and after exposure to hyperoxia (31 hours, see Materials and Methods) were assayed rapidly (hatched bars), reflecting the native activation state, or after pre-incubation for 10 minutes in the presence of $MgCl_2$, $H^{12}CO_3^-$, and 6-phosphogluconate, which promotes reactivation of inhibited enzyme (solid bars). Panel B) Table of values. Error bars represent the standard deviation of the three biological replicates, each with three technical replicates.

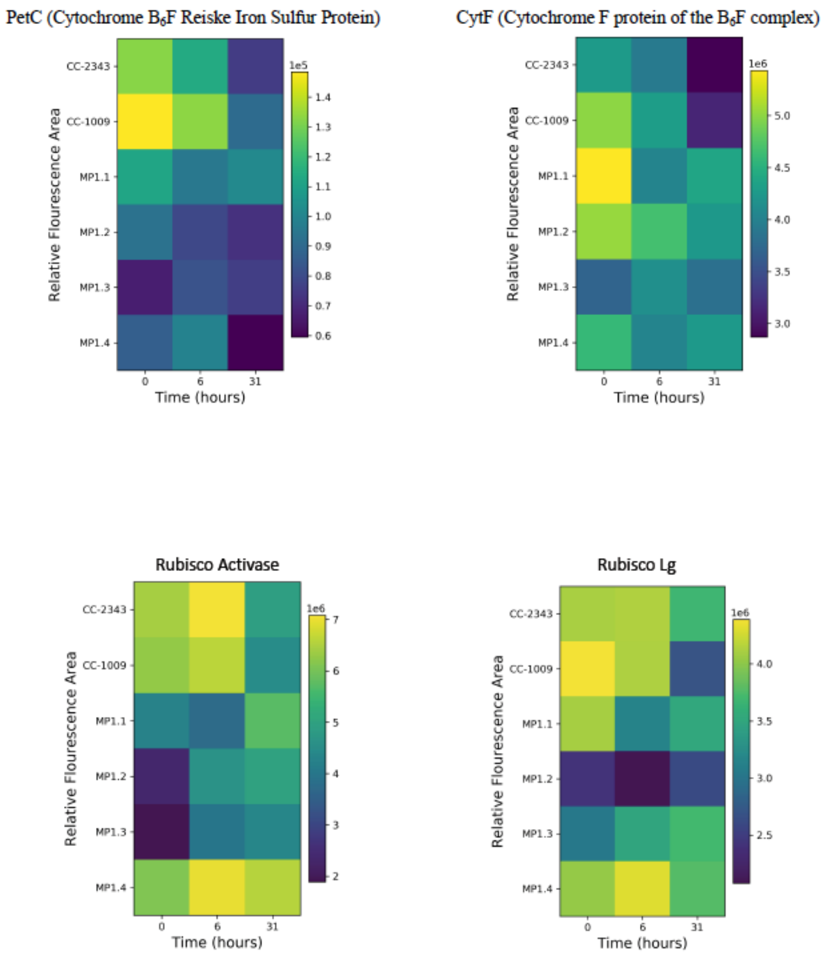


**Supplementary Material Figure 4**: Measured relative fluorescence areas of four different proteins involved in in the electron transport or the carboxylation reactions of photosynthesis. The proteins are CytF (Cytochrome F protein of the $B_6F$ complex); PetC (Cytochrome B6F Reiske Iron Sulfur Protein); Rubisco Activase; and Rubisco Lg (Rubisco Large Subunit). All were normalized to t=0. Unlike the pyrenoid formation, the abundance of none of these proteins altered in abundance in a manner consistent with tolerance to hyperoxia. The tolerant lines (CC-1009, c1_1, c1_2) do not show greater upregulation or these proteins in response to the stress.

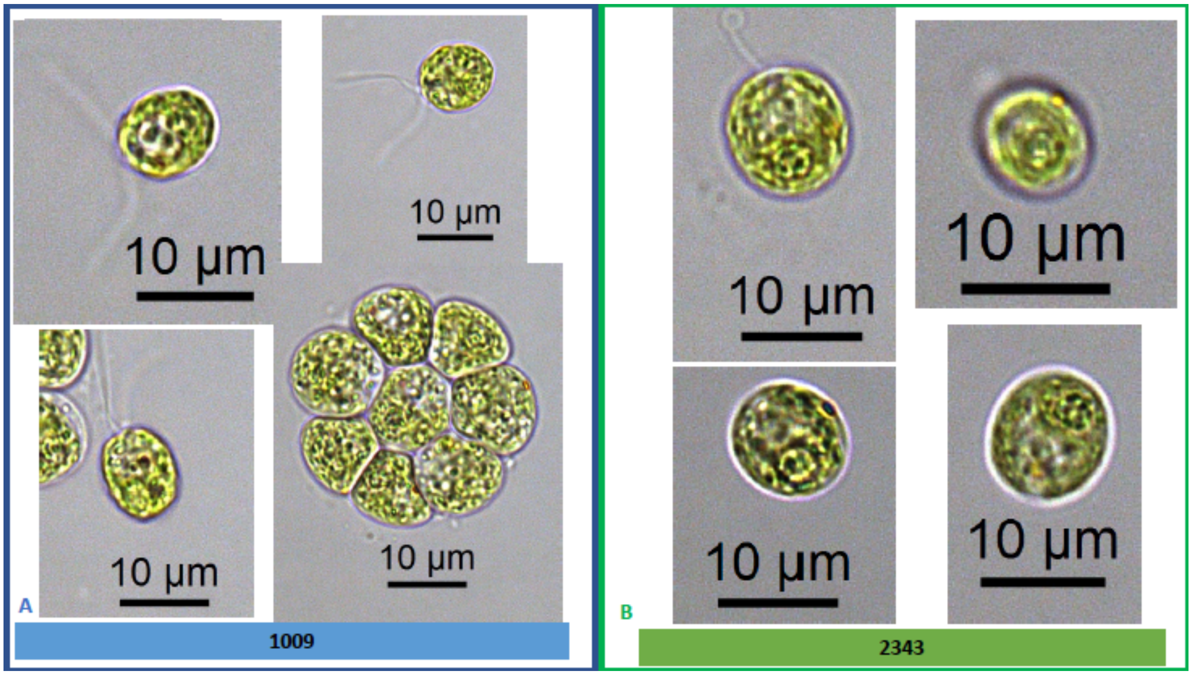


**Supplementary Material Figure 5**: Exemplary light microscopy images of *Chlamydomonas* strains (CC-1009 & CC-2343), while growing under saturating $CO_2$. Cells are growing with 5% $CO_2$ with 1 minute on/one minute off sparging, and 14:10 hour (light:dark) sinusoidal illumination with peak light intensity of 2000 μmoles $m^{-2}$ $s^{-1}$, in minimal 2NBH media. Cells here were viewed at at noon, at 2000 μmoles $m^{-2}$ $s^{-1}$. Sale bar = 2 μm.

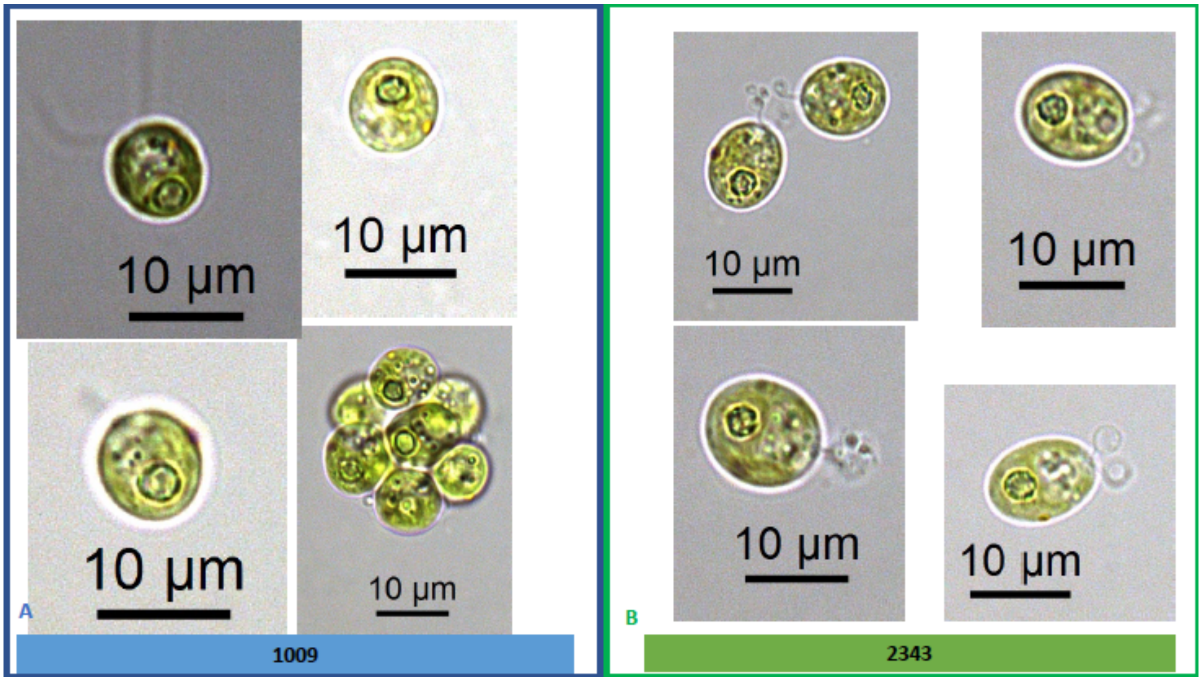


**Supplementary Material Figure 6**: Exemplary light microscopy images of *Chlamydomonas* strains (CC-1009 & CC-2343), after growing under saturating $CO_2$ and hyperoxia for 6 hours. Pyrenoid starch sheaths were clearly visible. Cells are sparged with 5% $CO_2$ and 95% $O_2$, with 1 minute on/one minute off sparging, and 14:10 hour (light:dark) sinusoidal illumination with peak light intensity of 2000 µmoles $m^{-2}$ $s^{-1}$, in minimal 2NBH media. Prior to switching the gas to hyperoxia (i.e. 95% $O_2$ and 5% $CO_2$) cells had been grown in steady state conditions. Cells here were viewed at noon, at 2000 µmoles $m^{-2}$ $s^{-1}$.

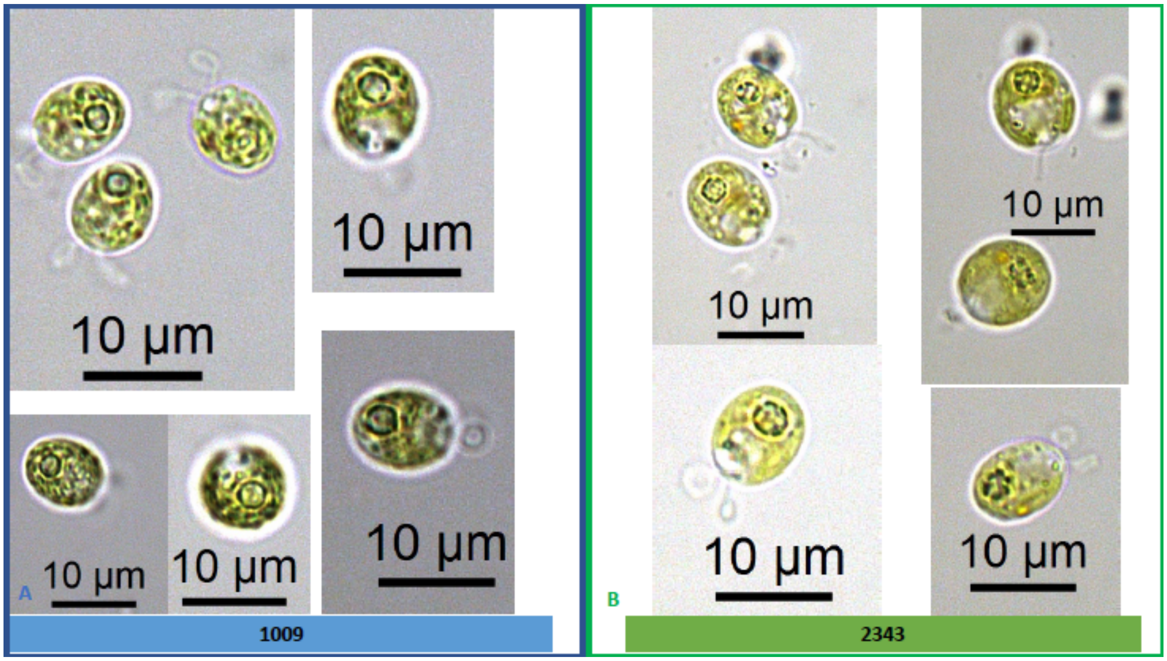


**Supplementary Material Figure 7**: Exemplary light microscopy images of *Chlamydomonas* strains (CC-1009 & CC-2343), after growing under saturating $CO_2$ and hyperoxia for 31 hours. Differences were observed between the pyrenoids of the cells in Panel A and Panel B, with the cells in Panel A showing more robust, continuous, sealed pyrenoids. Cells are sparged with 5% $CO_2$ and 95% $O_2$, with 1 minute on/one minute off sparging, and 14:10 hour (light:dark) sinusoidal illumination with peak light intensity of 2000 µmoles $m^{-2}$ $s^{-1}$, in minimal 2NBH media. Prior to switching the gas to hyperoxia (i.e. 95% $O_2$ and 5% $CO_2$) cells had been grown in steady state conditions. Cells here were viewed at noon, at 2000 µmoles $m^{-2}$ $s^{-1}$.

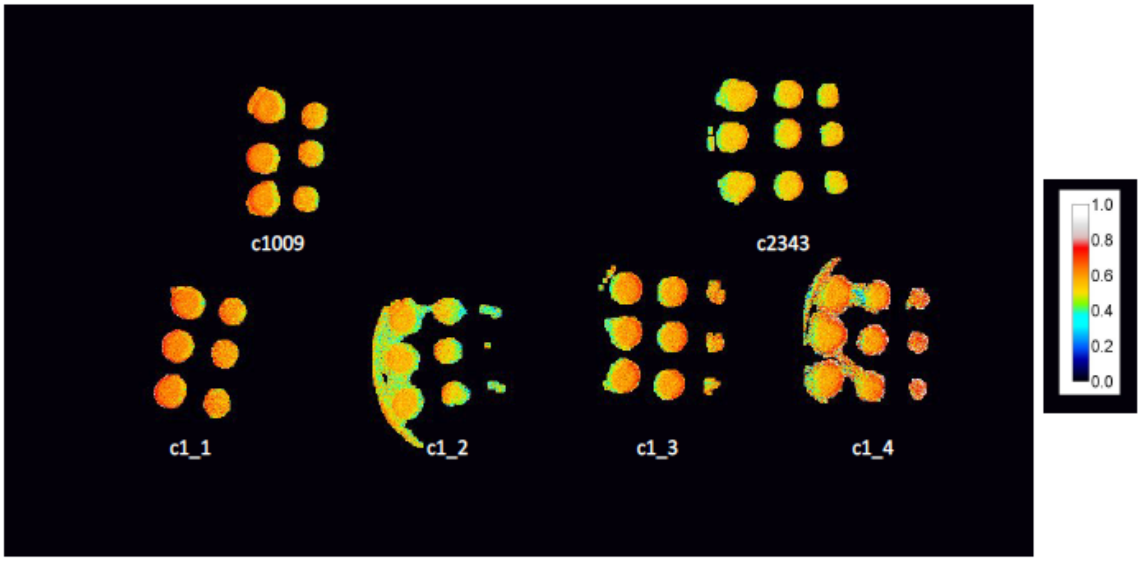


**Supplementary Material Figure 8**: $\Phi_{II}$ values of spotted plates of parents and tetrad after one week, with c1_3, and c1_4 not showing any obvious higher level of electron transport, despite higher growth rates. Measurements shown here were under 50 μM of PAR, though similar results were obtained up to 1500 μmoles $m^{-2}$ $s^{-2}$.

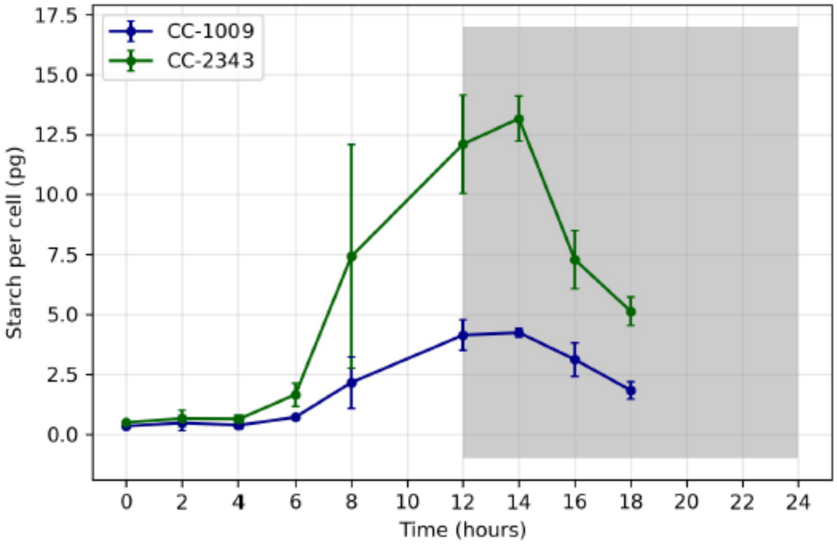


**Supplementary Material Figure 9**: Total starch accumulated (pg/cell) in CC-1009 and CC-2343 cultures grown at steady state conditions, with 5% $CO_2$ with 14:10 hour (light:dark) sinusoidal illumination with peak light intensity of 2000 µmoles $m^{-2}$ $s^{-1}$, in minimal 2NBH media . For TEM of steady state cells see Figure 6. Error bars represent the standard deviation of the three biological replicates.

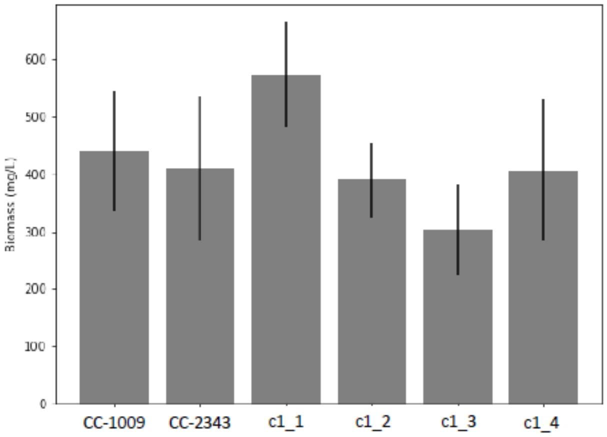


**Supplementary Material Figure 10**: Biomass measurement of CC-1009 and CC-2343 and tetrad progeny grown under 85 µmoles $m^{-2}$ $s^{-1}$ of light and low $CO_2$ in flasks for 4 days, after each flask was inoculated with $1x10^5$ cells/ml. Measurements were based on ash free dry weight of three different biological replicates for each strain. Error bars represent the standard deviation of three biological replicates. There was no statistical difference between the parents (CC-1009 and CC-2343) and any progeny. Though c1_1 did have higher productivity that two of the other progeny, all of the lines were able to grow under low $CO_2$ (i.e. none were high $CO_2$ requiring).

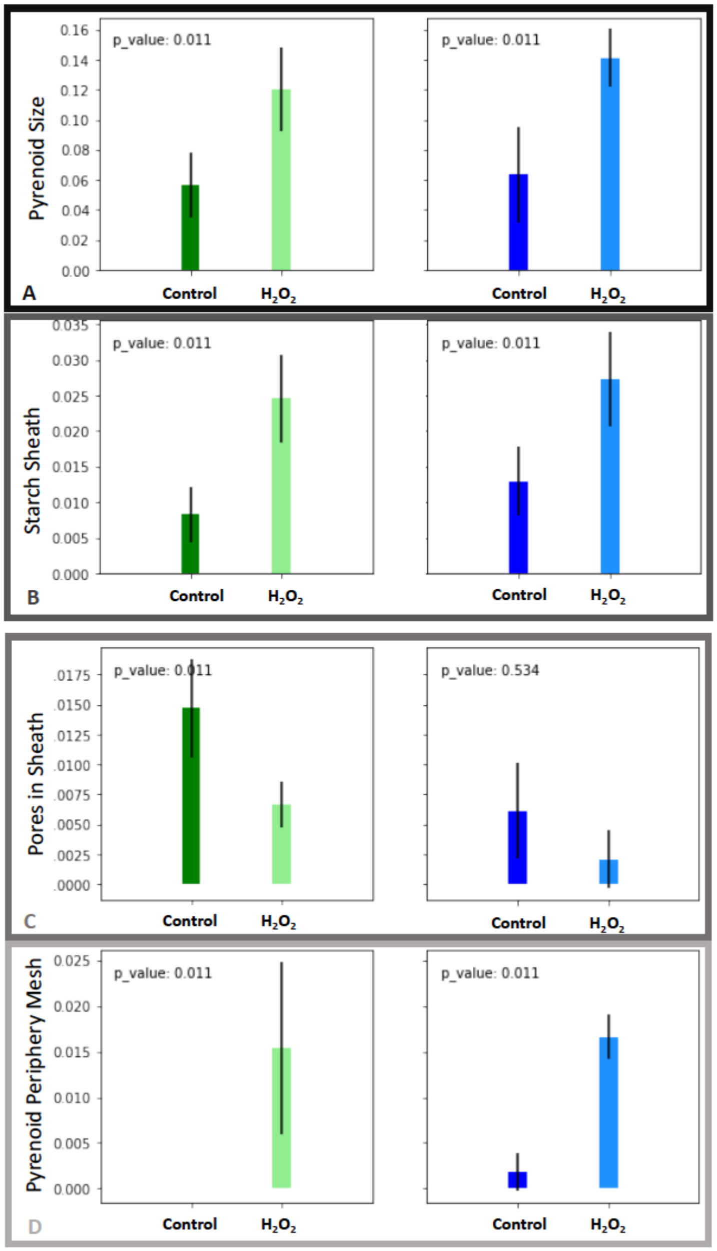


**Supplementary Material Figure 11**: Morphometric analysis of **CC-2343** and **CC-1009** cells, control (**green**, **blue**) and exposed to $H_2O_2$ (**light green**, **light blue**), normalized to cell size. In response to pre-treatment with $H_2O_2$, pyrenoid size (Panel A), and amount of starch in the sheath (Panel B) increased. Error bars represent the standard deviation. The addition of $H_2O_2$ also brought about the appearance of the pyrenoid periphery mesh, a matrix which appears to cement together the starch plates (Panel D). For CC-2343, which under control conditions had more gaps between the starch plates, the $H_2O_2$ also lead to a significant decline these gaps (Panel C).

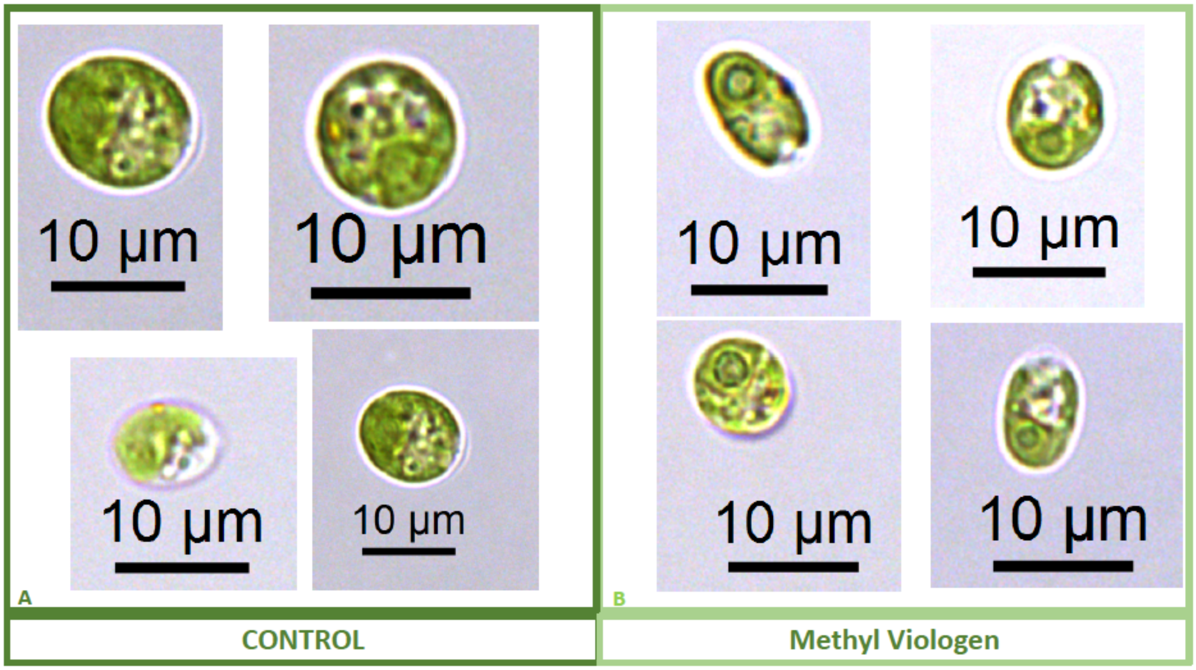


**Supplementary Material Figure 12**: Representative light microscopy images of CC-2343 control (Panel A) and treated (Panel B) cells, two hours after our sinusoidal light had turned on, with .1µM of methyl viologen, and then exposed to 6 hours of low light (~50 µmoles photons $m^{-2}$ $s^{-1}$) with saturating (5mM) bicarbonate in minimal 2NBH media.

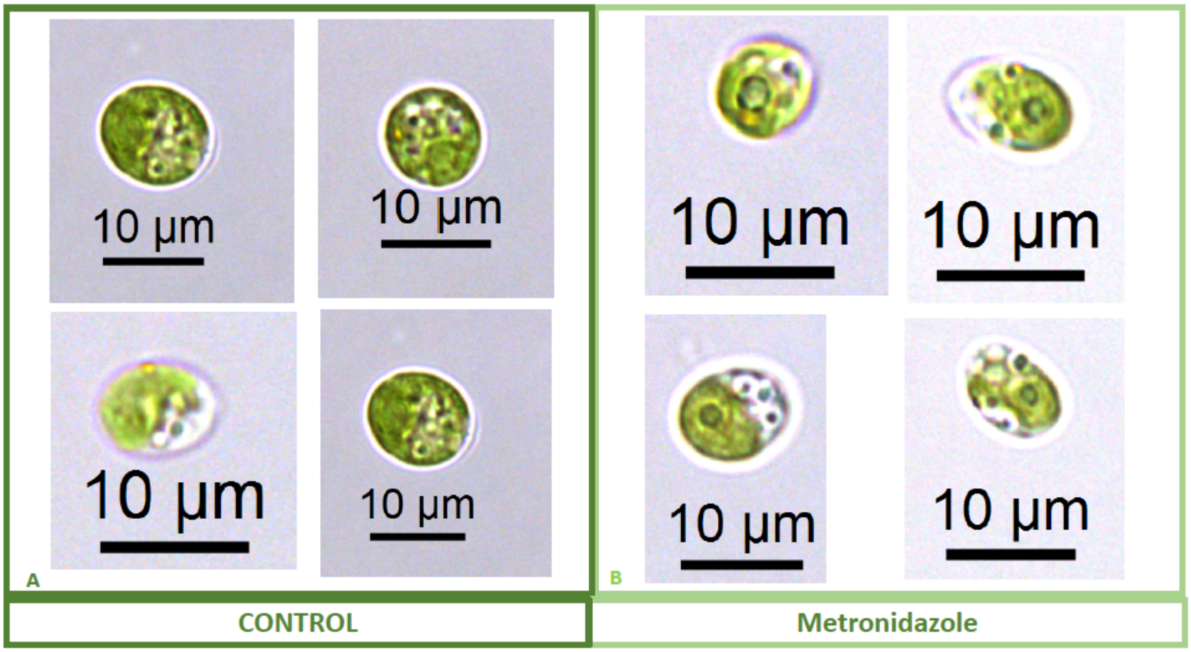


**Supplementary Material Figure 13**: Representative light microscopy images of CC-2343 control (Panel A) and treated (Panel B) cells, two hours after our sinusoidal light had turned on, with 4mM of metronidazole, and then exposed to 6 hours of low light (~50 µmoles photons $m^{-2}$ $s^{-1}$) with saturating (5mM) bicarbonate in minimal 2NBH media.

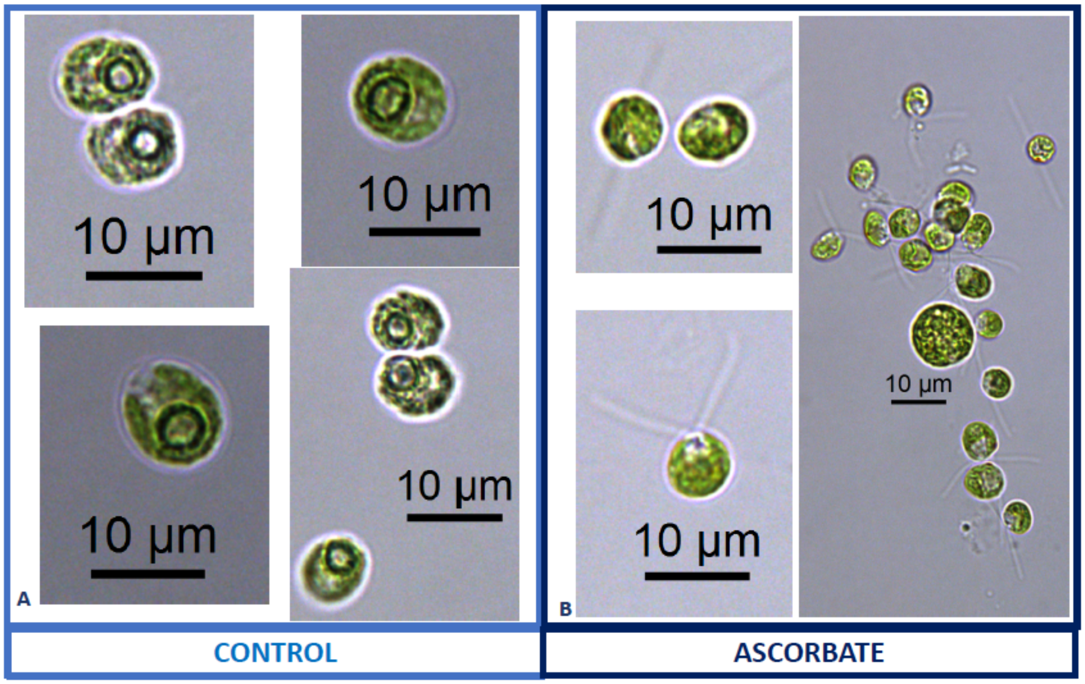


**Supplementary Material Figure 14**: Representative light microscopy images of CC-1009 control (Panel A) and 10mM ascorbate treated (Panel B) cells, grown for 12 hours while being bubbled with air (low $CO_2$) in minimal 2NBH media under ~85 µmoles photons $m^{-2}$ $s^{-1}$ .

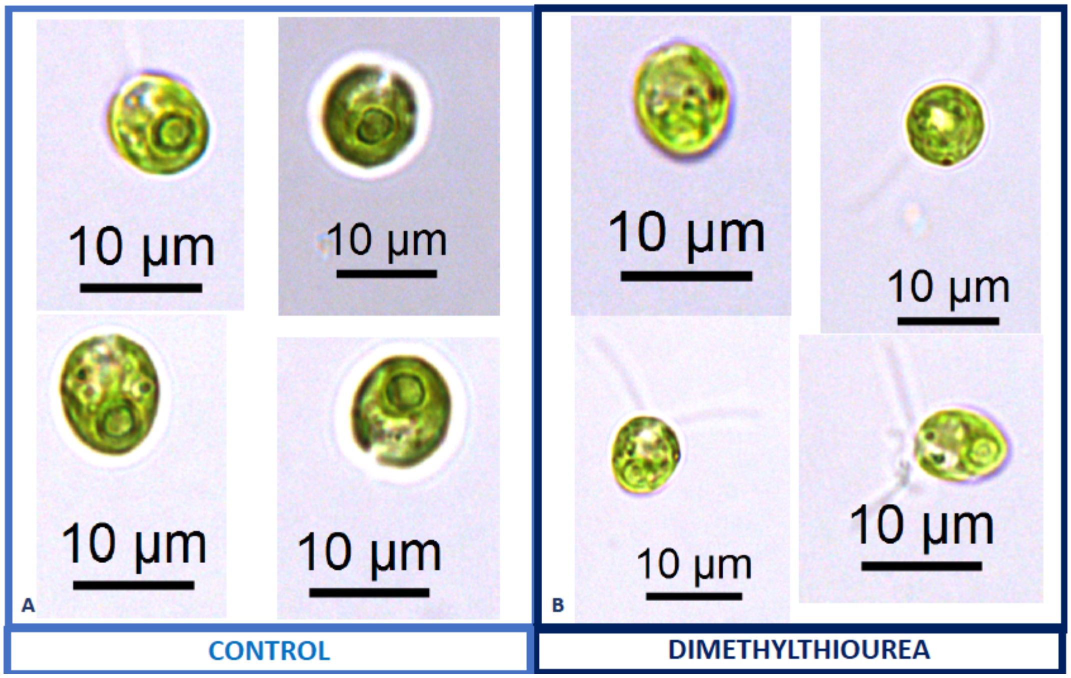


**Supplementary Material Figure 15**: Representative light microscopy images of CC-1009 control (Panel A) and 15mM dimethylthiourea treated (Panel B) cells, grown for 12 hours while being bubbled with air (low $CO_2$) in minimal 2NBH media under ~85 µmoles photons $m^{-2}$ $s^{-1}$ .

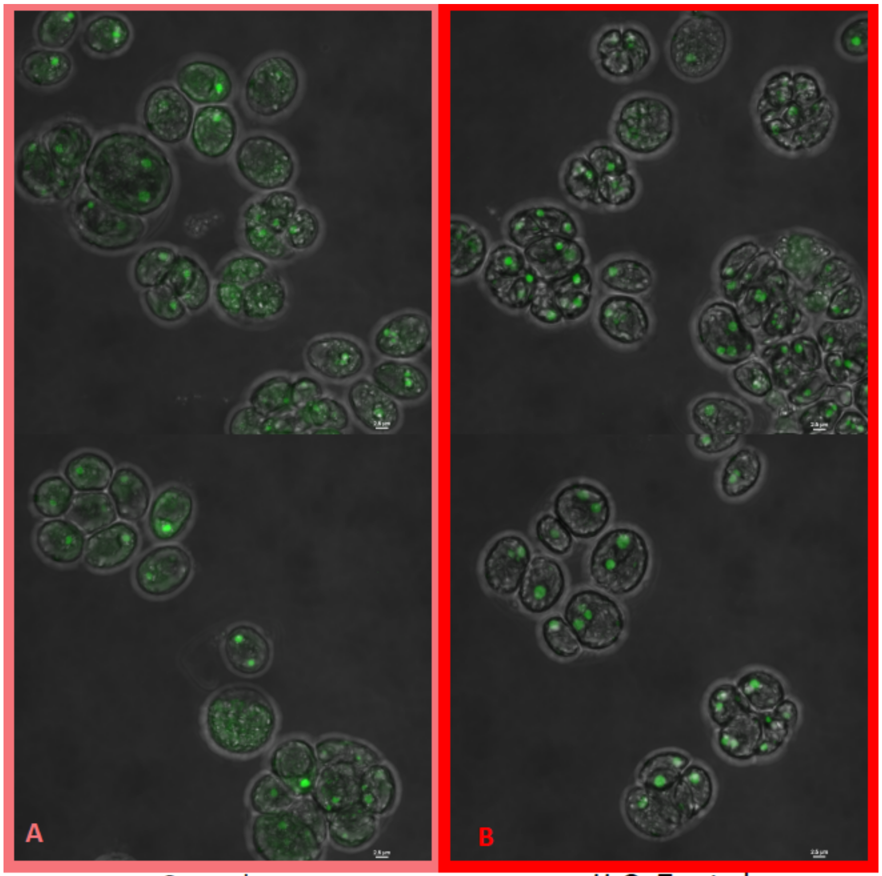


Control $H_2O_2$ Treated

**Supplementary Material Figure 16**: Localization of rubisco, control cells (Panel A) and cells exposed to 100µM $H_2O_2$, visualized by the Nikon A1 Confocal Laser Scanning Confocal Microscope confocal microscopy, in strain CC-5357, containing at RBCS1-Venus. Scale bar = 2 µm. These images, along with those with the Olympus (Figure 10), indicate a clear change in rubisco localization, rather the simply a change in rubisco amount. Since confocal microscopy is designed to acquire a very thin optical section through the thickness of a single cell, it can be difficult to use confocal fluorescence microscopy to accurately measure total protein content within the 3-dimensional volume of a single cell. Acquisition of several images through the thickness of the cell may actually over estimate or under estimate the total fluorescence intensity, depending on the the Z-step increment. However, when comparing these images of Venus Fluorescent Protein-labeled rubisco within *Chlamydomonas* cells acquired using the Nikon A1 confocal microscope, the fluorescence intensity of the pyrenoid matrix of most cells under control conditions (5 mM bicarbonate, no $H_2O_2$ treatment) was similar to or dimmer than the fluorescence intensity of the pyrenoid matrix within cells treated for six hours with 100 µM $H_2O_2$. In contrast, the fluorescence intensity of the Venus Fluorescent Protein-labeled rubisco located outside of the pyrenoid matrix was noticeably (measurably) higher in cells under control conditions compared to $H_2O_2$-treated cells. If the decrease in Venus Fluorescent Protein-labeled rubisco fluorescence located outside of the pyrenoid matrix in $H_2O_2$-treated cells was due to an overall decrease in rubisco production within the cell and not due to delocalization of the protein from the pyrenoid to the cytoplasm of the cell, then fluorescence intensity of the pyrenoid within $H_2O_2$-treated cells should show a comparable decrease in fluorescence intensity, which is not seen.

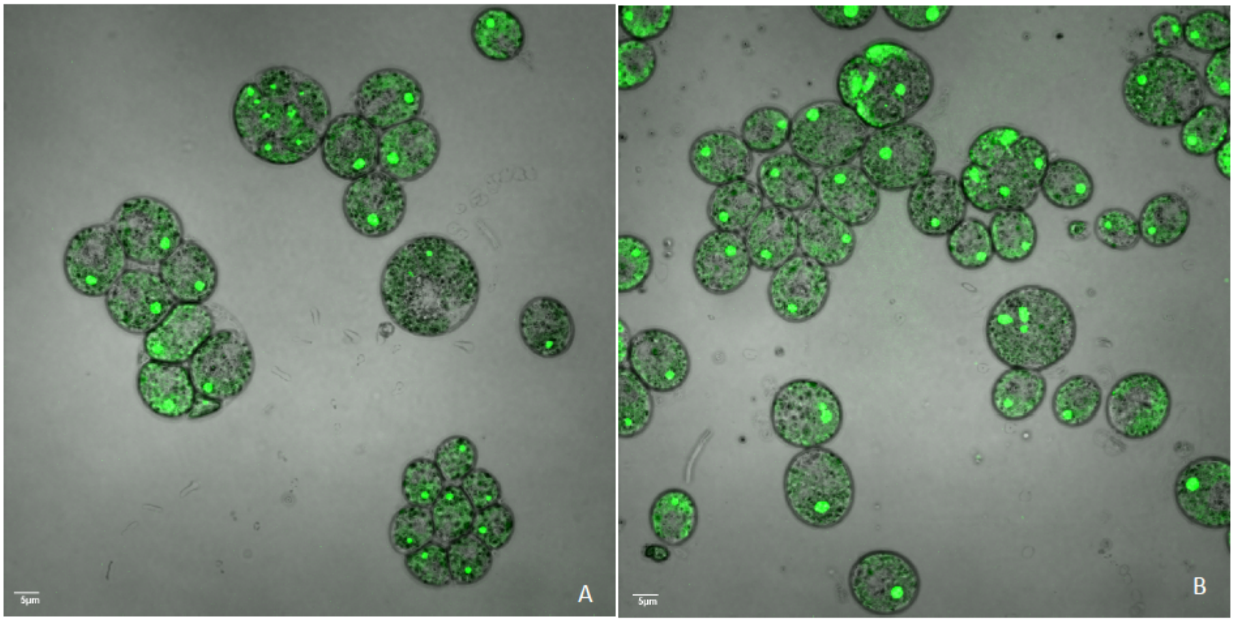


**Supplementary Material Figure 17**: Confocal microscopy of liquid TAP grown CC-5357, which has a Venus labeled RBCS1, treated with (Panel A) and without $H_2O_2$ (Panel B), showing no change in localization of rubisco.

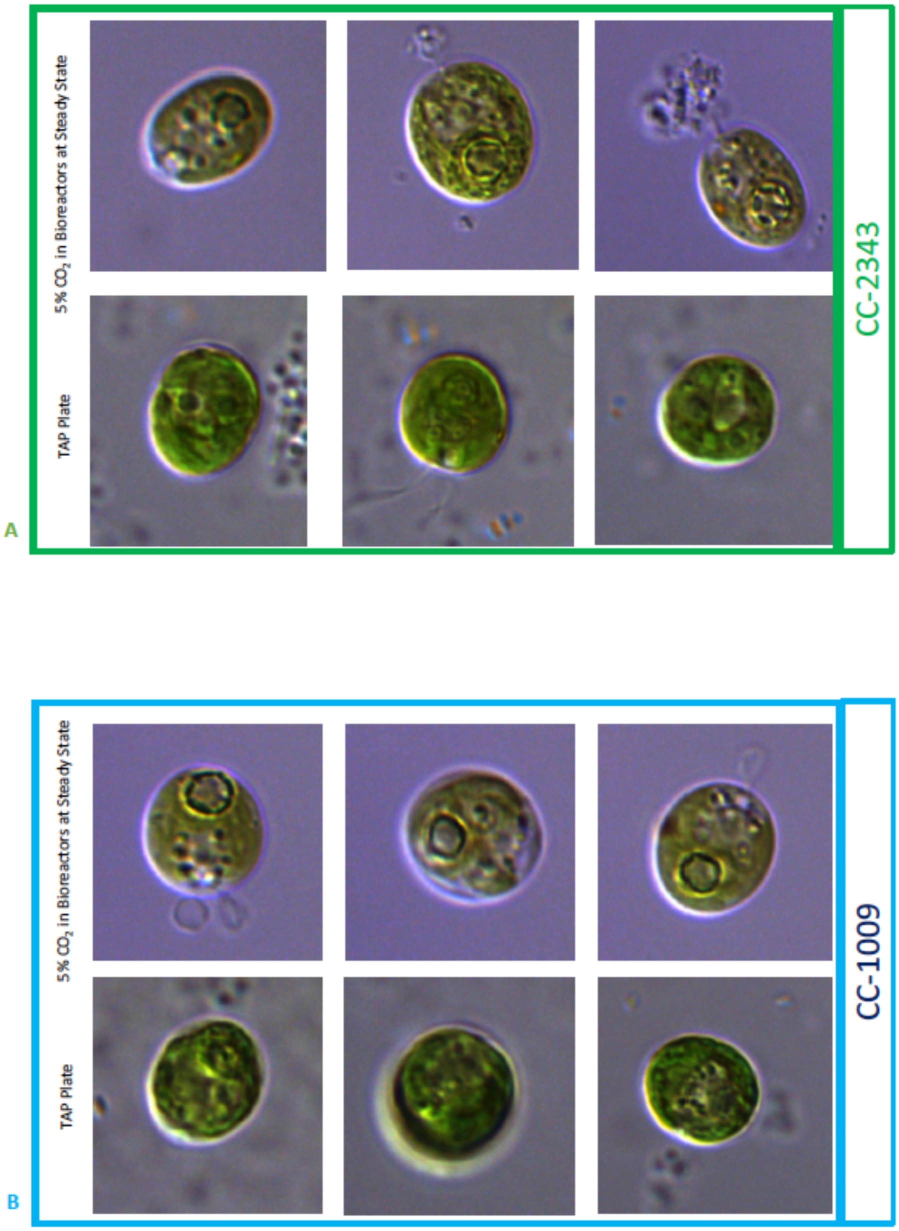


**Supplementary Material Figure 18**: Comparison of cells grown under steady state in liquid media (with 5% $CO_2$) versus cells grown on the surface of a TAP plate. Both CC-2343 (Panel A) and CC-1009 (Panel B) showed losses of the starch sheath when grown on a TAP plate.

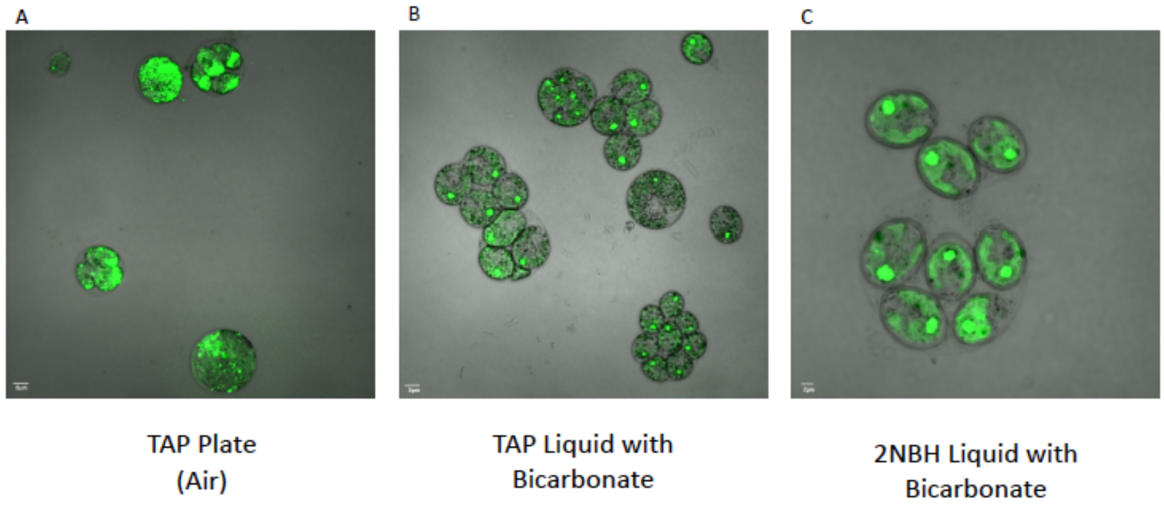


**Supplementary Material Figure 19**: Comparison of CC-5357, with a fluorescently-tagged rubisco, grown at various conditions. Panels A & B, Scale bar = 5 µm, Panels C = 2 µm.

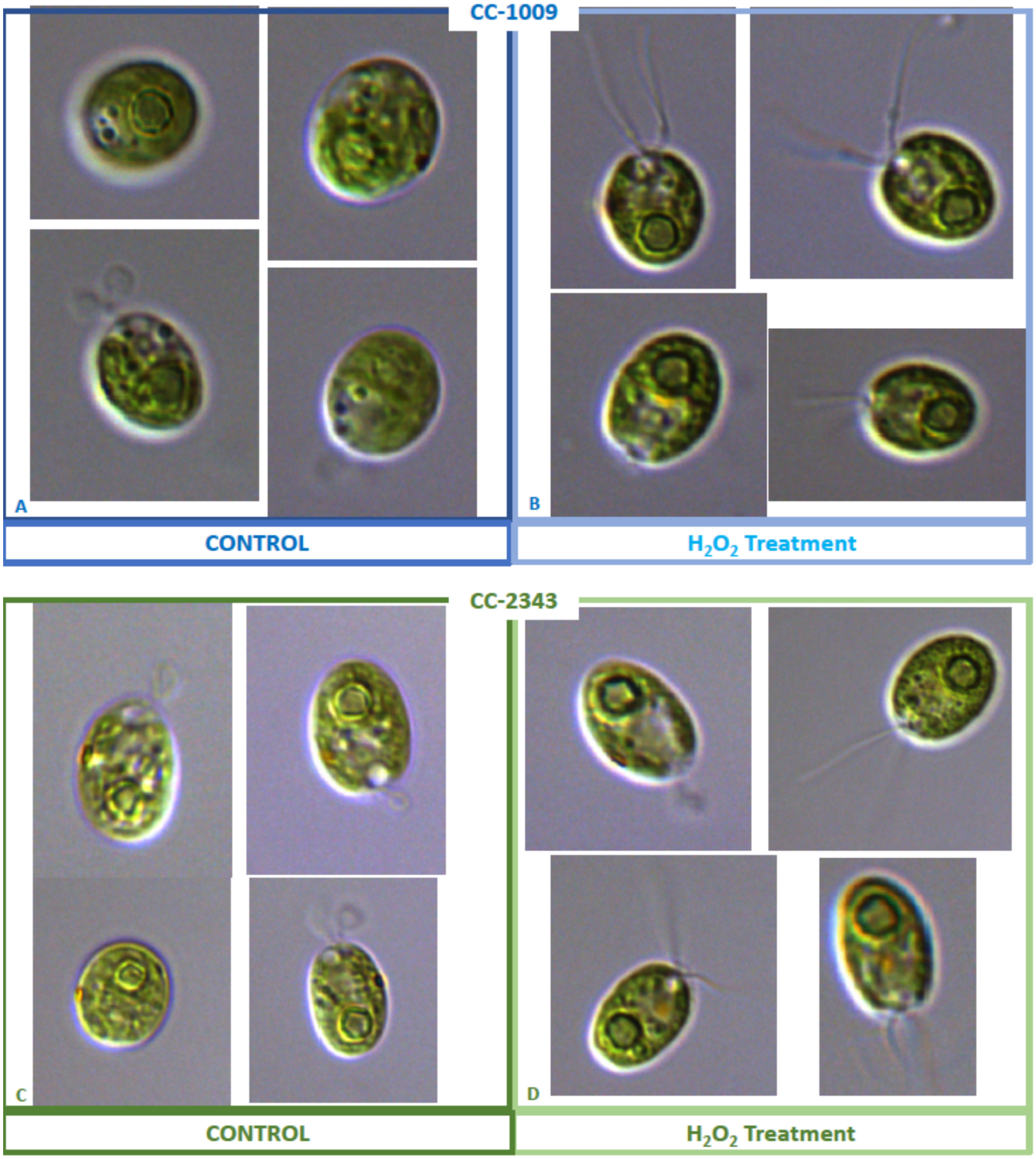


**Supplementary Material Figure 20**: Representative light microscopy images of CC-1009 (Panels A & B) and CC-2343 (Panels C & D) control and cells treated, two hours after our sinusoidal light had turned on, with 100 µM of $H_2O_2$, and then exposed to 6 hours of low light (~50 µmoles photons $m^{-2}$ $s^{-1}$) with saturating (5mM) bicarbonate in minimal 2NBH media.

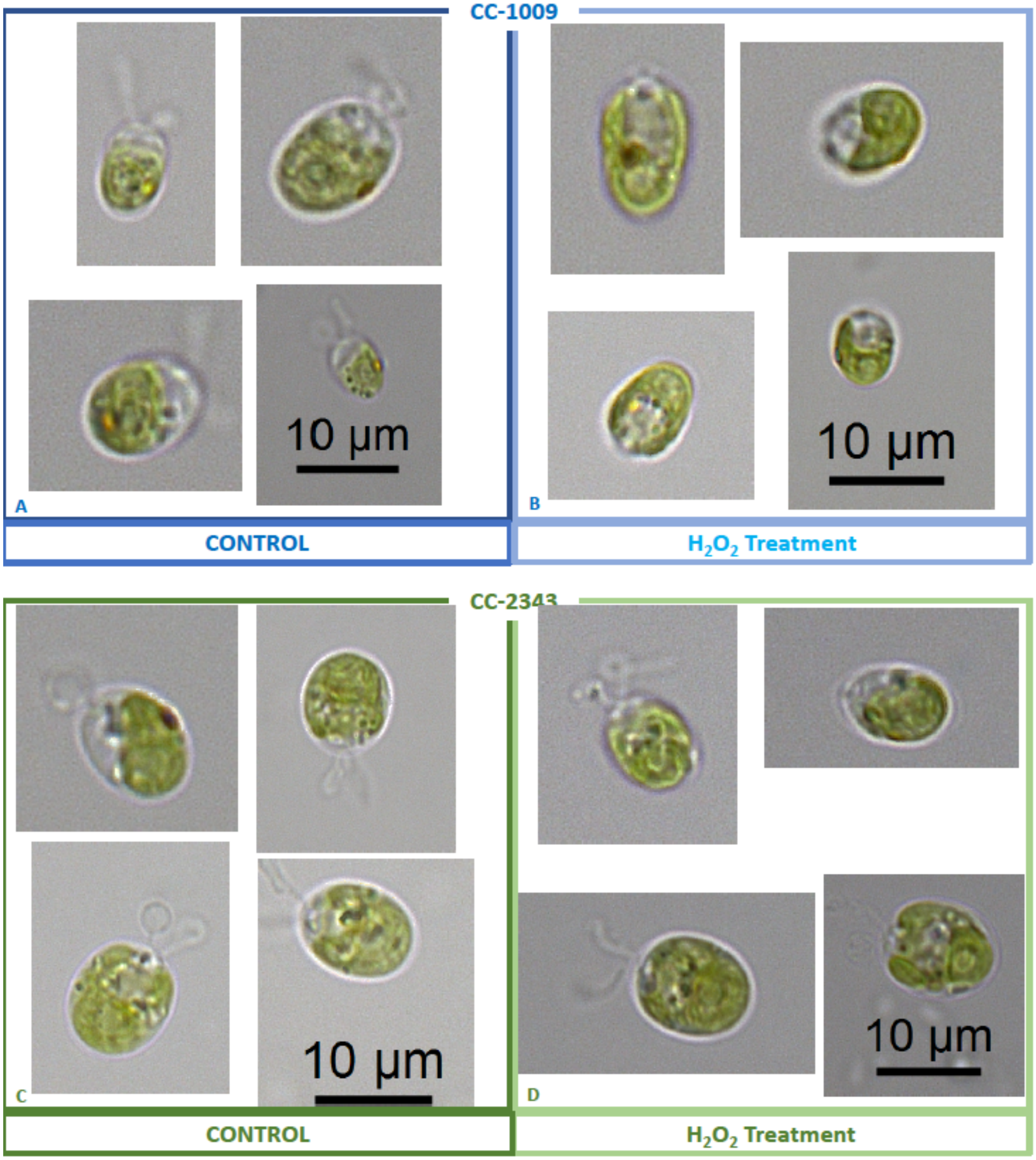


**Supplementary Material Figure 21**: Representative light microscopy images of CC-1009 (Panels A & B) and CC-2343 (Panels C & D) control and cells treated, the light remaining off, with 100 µM of $H_2O_2$, and kept at 6 hours of dark with saturating (5mM) bicarbonate in minimal 2NBH media.
